# Nutritional inter-dependencies and a carbazole-dioxygenase are key elements of a bacterial consortium relying on a *Sphingomonas* for the degradation of the fungicide thiabendazole

**DOI:** 10.1101/2020.03.30.015693

**Authors:** Vasileiadis Sotirios, Perruchon Chiara, Scheer Benjamin, Adrian Lorenz, Steinbach Nicole, Trevisan Marco, Plaza-Bolaños Patricia, Agüera Ana, Chatzinotas Antonis, Karpouzas G Dimitrios

## Abstract

**Background:** Thiabendazole (TBZ), is a benzimidazole fungicide and anthelminthic whose high persistence and toxicity pose a serious environmental threat. In our quest for environmental mitigation we previously isolated the first TBZ-degrading bacterial consortium and provided preliminary evidence for its composition and the degrading role of a *Sphingomonas*. Here, we employed a multi-omic approach combined with DNA-stable isotope probing (SIP) to determine the genetic make-up of the key consortium members, to disentangle nutritional and metabolic interdependencies, to identify the transformation pathway of TBZ and to understand the genetic network driving its transformation.

**Results:** Time-series SIP in combination with amplicon sequencing analysis verified the key role of *Sphingomonas* in TBZ degradation by assimilating over 80% of the ^13^C-labelled phenyl moiety of TBZ. Non-target mass spectroscopy (MS) analysis showed the accumulation of thiazole-4-carboxamidine as a single dead-end transformation product and no phenyl-containing derivative, in line with the phenyl moiety assimilation in the SIP analysis. Time series metagenomic analysis of the consortium supplemented with TBZ or succinate led to the assembly of 18 metagenome-assembled genomes (MAGs) with >80% completeness, six (*Sphingomonas* 3X21F, *γ-Proteobacterium* 34A, *Bradyrhizobiaceae* 9B and *Hydrogenophaga* 19A, 13A, and 23F) being dominant. Meta-transcriptomic and -proteomic analysis suggested that *Sphingomonas* mobilize a carbazole dioxygenase (*car*) operon during the initial cleavage of TBZ to thiazole-4-carboxamidine and catechol, the latter is further transformed by enzymes encoded in a catechol *ortho*-cleavage (*cat*) operon; both operons being up-regulated during TBZ degradation. Computational docking analysis of the terminal oxygenase component of *car*, CarAa, showed high affinity to TBZ, comparable to carbazole, reinforcing its high potency for TBZ transformation. These results suggest no interactions between consortium members in TBZ transformation, performed solely by *Sphingomonas*. In contrast, gene expression network analysis revealed strong interactions between *Sphingomonas* MAG 3X12F and *Hydrogenophaga* MAG 23F, with *Hydrogenophaga* activating its cobalamin biosynthetic pathway and *Sphingomonas* its cobalamin salvage pathway along TBZ degradation.

**Conclusions:** Our findings suggest interactions between consortium members which align with the “black queen hypothesis”: *Sphingomonas* detoxifies TBZ, releasing consortium members by a toxicant; in return for this, *Hydrogenophaga* 23F provides cobalamin to the auxotrophic *Sphingomonas*.

## Background

Thiabendazole (TBZ) is a benzimidazole compound which is used as a post-harvest fungicide to control fungal infestations on fruits during storage [1] and as a broad spectrum anthelminthic to control endoparasites in livestock farming [2]. It acts by binding to tubulin monomers inhibiting the polymerization of microtubules and, thus, cell growth [3–5]. TBZ has been identified as a common contaminant of natural water resources in citrus producing areas [6, 7] threatening the integrity of water ecosystems due to its high aquatic toxicity (i.e. NOEC fish = 12 μg L^−1^) [8]. We recently reported TBZ concentration levels in soils adjacent to fruit packaging plants of up to 12,000 mg kg^−1^ resulting in a depleted bacterial diversity [9]. The extensive environmental contamination by TBZ is the result of its high persistence (DT50 (soil - aerobic)> 365 days) [10] and the lack of implemented methods for the treatment of TBZ-contaminated agro-industrial effluents (despite the implementation of relevant EC legislation) [11]. The problem is further exemplified by the limited capacity of microbial communities in municipal wastewater treatment plants to remove recalcitrant chemicals like TBZ [12]. Instead municipal wastewater treatment plants act as point sources for the contamination of receiving water bodies [12] and agricultural soils [13]. In the latter case the application of biosolids, derived from a TBZ-containing wastewater treatment plant, as fertilizers resulted in the persistence of 83% of the initially applied TBZ after 3 years [13].

Bacterial specialists that have the capacity to degrade TBZ could be invaluable in bioaugmentation and biodepuration strategies to avert its environmental impact. In this context, we recently enriched from a heavily TBZ contaminated soil the first bacterial consortium capable of degrading and detoxifying TBZ while using it as the sole carbon source [14, 15]. The consortium, which was dominated by different α-, β- and *γ-Proteobacteria*, was stable in its composition and its degrading efficiency. Preliminary assays (i.e. stable isotope probing combined with denaturant gradient gel electrophoresis (SIP-DGGE) analysis) pointed to a *Sphingomonas* as the key degrader of TBZ [14], with the roles of the other consortium members remaining unknown. No single TBZ-degrading bacterial strain was isolated from the consortium despite our copious attempts with different media and solidifying agents, indicating underlying interactions of the key degrader with the other members of the consortium. The coherence of pollutant-degrading microbial consortia goes beyond simple collaborative transformation of the target pollutants [16, 17] and involves syntrophic and cross-feeding relationships on biomolecules like amino acids and vitamins [18–20]. Comparative genomic analyses suggested an evolutionary drift in bacterial genomes towards auxotrophic lifestyles on energetically costly amino acids and cofactors like B12 [21, 22], shaping microbial communities in various environments and the human gut [23].

A prerequisite for the biotechnological exploitation of microbial consortia is to disentangle the roles of keystone members, degraders and suppliers/feeders on key nutrients, whose presence guarantees consortium coherence [24, 25]. In the current study we seek answers to the following questions: (i) Is *Sphingomonas* the sole member of the consortium involved in the transformation of TBZ or are there transformation interdependencies driving the detoxification the fungicide? (ii) Which are the main transformation products of TBZ? (iii) Which is the role of other members of the consortium, with respect to TBZ degradation, and does its stable composition infer established roles amongst them? (iv) Which are the genes and enzymes involved in the transformation process? To answer these questions we employed, in time series experiments, an integrated, multi-omics approach (meta-genomics/transcriptomics/proteomics) combined with non-target mass spectroscopy (MS) analysis and SIP-based amplicon sequencing, with the latter enabling the confident identification of the key degrading members of the consortium.

## Methods

### Microbial consortium growth conditions

The TBZ-degrading bacterial consortium was routinely grown at 27°C in a minimal salts medium (MSMN) supplemented with TBZ (25 mg l^−1^) as the sole carbon source [14] (see the supporting information - SI - for media composition).

### TBZ degradation assays

#### Experiment 1 - SIP analysis

A first degradation assay was performed to identify the members of the consortium involved in the transformation of TBZ via DNA-SIP based amplicon sequencing analysis. Triplicate 30-ml cultures of the consortium were supplemented with 25 mg l^−1^ of unlabelled (^12^C) or ^13^C-TBZ labelled uniformly in its phenyl moiety (Clearsynth®, Mumbai, India). Triplicate flasks of MSMN supplemented with unlabelled TBZ but not inoculated with the consortium were co-incubated as abiotic controls. Aliquots of the cultures (0.5 ml) were removed at 36, 72, 117 and 141 h, (corresponding to 10%, 30%, 100% degradation of TBZ and 24 h after its complete degradation, Figure 1A) to determine: (i) TBZ degradation via HPLC analysis as described previously [14] and (ii) community composition via DNA extraction and amplicon sequencing as described below.

**Figure 1.**
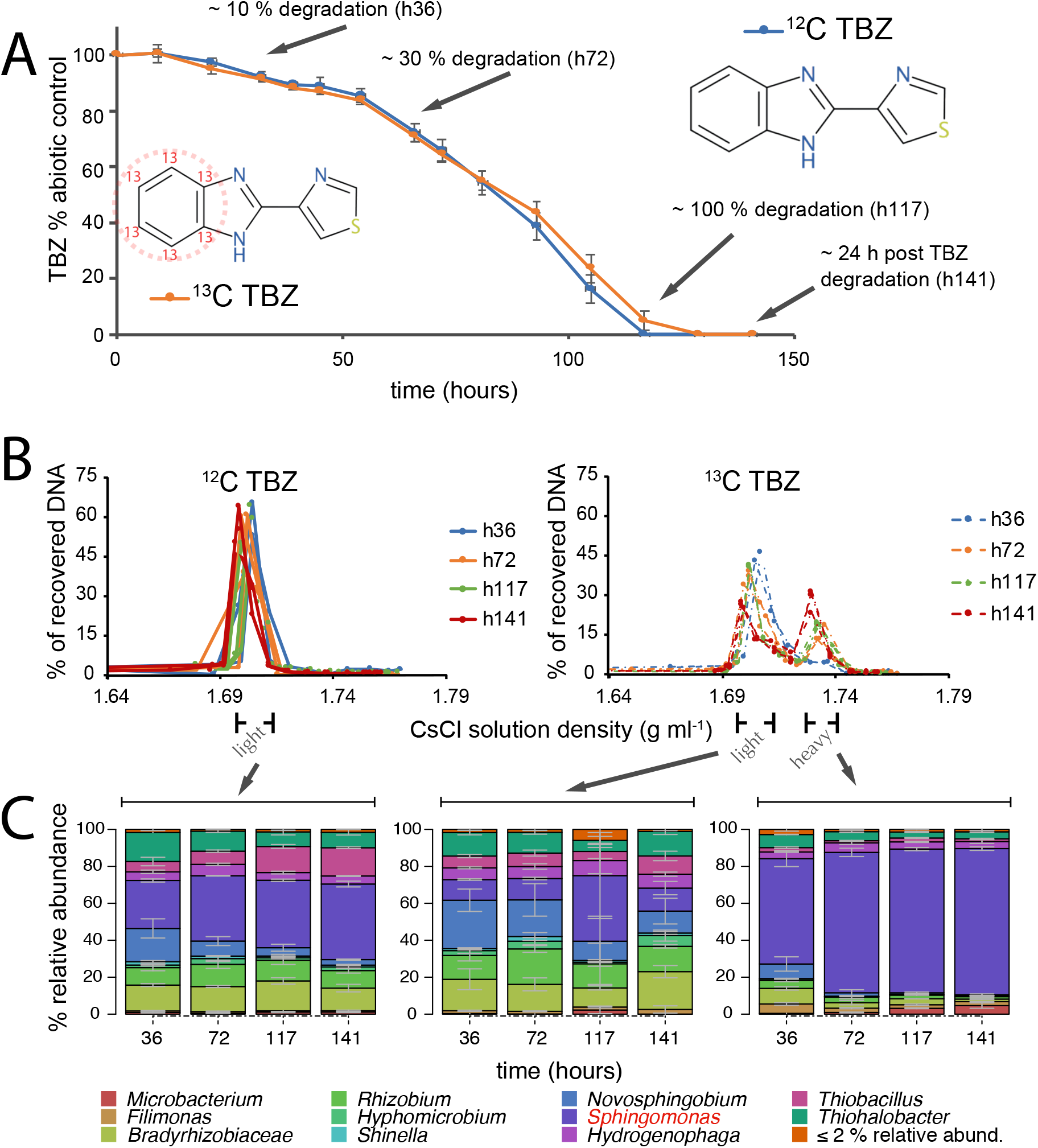
(A) The degradation patterns of unlabelled (^12^C, blue line) and ^13^C-labelled TBZ (red line) in MSMN inoculated with the bacterial consortium. Data are presented as % degradation relatively to the non-inoculated abiotic control. Each value is the mean of three replicates + the standard deviation. Arrows indicate the time points (and % degradation of TBZ) where amplicon sequencing analysis was implemented; (B) Recovery of DNA after density gradient centrifugation of sample replicates retrieved from the unlabelled TBZ supplemented culture (left) and the ^13^C-labelled-TBZ grown cultures with the light (corresponding to the unlabelled TBZ sample peak CsCl solution densities of 1.69-1.72 g ml^−1^) and the heavy DNA fraction (corresponding to CsCl solution densities of 1.72-1.75 g ml^−1^) separated along the CsCl density gradient (right panel); (C) The composition of the bacterial consortium (determined via 16S rRNA gene amplicon sequencing) in the different DNA fractions (unlabelled, light and heavy fraction of ^13^C-labelled TBZ) described above, with stacked bars presenting the mean relative abundances of the OTUs among triplicates, and error bars showing the standard deviations (taxa with up to 2 % relative abundance in all samples were grouped together).

#### Experiment 2 - multi-omic analyses

A second degradation assay was employed to disentangle metabolic interactions between consortium members and to identify, *via* a multi-omic approach, the key genes/enzymes driving the transformation of TBZ. Triplicate cultures of the consortium were amended either with 25 mg l^−1^ TBZ (125 μM) or 37 mg l^−1^ of succinate (SUC; 314 μM) as the sole carbon source (with a carbon concentration of 15 μg ml^−1^ in each case). Parallel triplicate abiotic controls as described above were also included. Aliquots (0.5 to 4 ml depending on the type of measurement employed) were removed from cultures at multiple time points along the degradation of TBZ and used for DNA/RNA/protein extraction and downstream metagenome binning, transcriptomic, proteomic and non-target MS analysis for TBZ transformation product detection.

### Nucleic acids extraction and quantification

DNA and RNA were extracted from bacterial cell pellets with the NucleoSpin^®^ Tissue and RNA kits, respectively (Macherey-Nagel & Co, Düren, Germany). Nucleic acid extracts were quantified with the Quant-iT^™^ HS ds-DNA assay kit and the Quanti-iT^™^ RNA HS kit with a Qubit^™^ fluorometer (Invitrogen, USA).

### SIP analysis

DNA extracts from the ^13^C-labelled TBZ-treated culture were separated into ^13^C-labelled and unlabelled fractions according to their buoyant density in a CsCl gradient established by ultracentrifugation at 167,000 *x g* for 36 hours at 20 °C [26]. DNA was extracted from the CsCl gradient buffer by glycogel/PEG precipitation and used for subsequent 16S rRNA gene diversity analysis as described further on and detailed in the SI.

### Amplicon sequencing analysis

The composition of the bacterial consortium in experiment 1 (SIP), was determined via multiplex sequencing of PCR amplicons of the V4 hypervariable region of the 16S rRNA gene according to our in-house protocol [27, 28] using primers 515F/806R [29, 30] as described in details in the SI. The sample-wise-demultiplexed/quality-controlled read pairs were used for reconstructing the amplicon sequences which were processed with the Lotus v1.58 suit [31] for generating the 97% sequence identity operational taxonomic unit (OTU) matrices and obtaining their taxonomic classifications. The β-diversity analysis was performed with the Entropart v1.4-7 [32] and the Vegan v2.5-5 [33] R v3.6.0 software [34] packages (more details are available in the SI).

### Metagenome assembly, contig binning, annotation and synteny analysis

Metagenome assembly within the context of the multi-omics experiment 2, was performed using the sequencing data of five shotgun libraries over three sequencing runs using both second (Illumina) and third generation (Pacific Biosciences) sequencing approaches as described in detail in the SI. The use of both second and third generation sequencing approaches along with the choice of samples of the consortium for DNA extraction at varying experimental conditions (assuring for differential genomic coverage by the sequencing reads), aimed at achieving a robust assembly of the metagenome and extracting metagenome assembled genomes (MAGs) as proposed previously [35]. Sequences were quality controlled and the devoted sequencing effort was assessed with Nonpareil v3.301 [36, 37]. Hybrid assembly was performed with Mira v5. 1 [38] and Megahit v1.1.3 [39] as described in the SI. Metawatt v3.5.3 [40] was used for obtaining MAGs which were further classified and quality assessed with MiGA against its registered NCBI genome collection [41]. Annotation of the sequences was performed with Prokka v1.12 [42], enriched with the aromatic hydrocarbon degradation AromaDeg [43] and the mobile genetic elements ALCME v0.4 [44] protein databases. The predicted open reading frames (ORFs) were also compared against the SEED database [45] with Rapsearch v2.22 [46]. BLAST was used for identifying molecular anchors during comparative genomics and the GenoPlotR v0.8.9 [47] R package was used for generating the associated plots.

### RNA sequencing analysis

Samples from experiment 2 obtained at 57, 73, and 109 hours post inoculation (hpi), corresponding to 40% degradation, 100% degradation and 36 h after 100% degradation of TBZ, were collected from TBZ amended bacterial cultures (and from the succinate amended cultures) for transcriptomic analysis via shotgun sequencing in Illumina Hiseq 2×250 bp rapid mode. Quality controlled sequences obtained were mapped against the reference metagenome assembly sequence with STAR v020201 [48], while transcript counts were predicted with HTSeq v0.9.1 [49]. Differential expression analysis was performed using the trimmed mean of M-values (TMM) normalization approach [50] with edgeR v3.14.0 [51] and hypotheses were tested with the negative binomial models and the generalized linear model quasi likelihood F-test [52]. Associated multivariate analysis and modeling was performed with the Vegan R package, while Spearman correlation tests (p ≥ 0.5; Benjamini-Hochberg adjusted P-value ≤ 0.05) followed by network analysis and associated substructure [53] identification methods were used for identifying transcript memberships (see SI). Network and the keystoneness [54] analyses were performed with Igraph v1.0.1 [55].

### Metaproteomic analysis

Lysed cells from the same samples used for RNA sequencing were treated with dithiothreitol as reducing agent and iodoacetamide to break and prevent the reformation of disulfide bonds respectively.

The resulting protein extracts were digested with trypsin, and the samples were desalted with ZipTips as described in detail in the SI. Peptide mixtures were analyzed by nanoLC-MS/MS using an Orbitrap Fusion mass spectrometer (ThermoFisher Scientific, Waltham, MA, USA). Protein identification was performed as described previously [56] with Proteome Discoverer v2.2 (ThermoFisher Scientific, Waltham, MA, USA) using SequestHT to search against the consortium metagenome translations of the predicted ORFs. Label-free quantification of peptides was done with the Minora node implemented in Proteome Discoverer. The abundances of confidently predicted proteins (false discovery rate below 1% as determined with the Percolator node) were analysed similarly with the RNA sequencing data (see SI).

### Non-target MS analysis of TBZ transformation products

Samples from experiment 2 collected from different time points along the degradation of TBZ were lysed by four repeats of sonication at 80 kHz for 30 s. They were then filtered through 0.22-μm PTFE syringe filters and aliquots of 450 μl were diluted with 25 μl of acetonitrile and 25 μl of a ^13^C-caffeine (internal) standard solution (Sigma-Aldrich, Steinheim, Germany). These were injected in a liquid chromatography coupled with a quadrupole-time-of-flight mass analyser (LC-QTOF-MS) with an Agilent 1260 Infinity system (Agilent Technologies, Foster City, CA, USA) connected to a Triple TOF 5600+ (Sciex Instruments, Foster City, CA, USA). The chromatographic separation was performed using a SB-C18 analytical column (3 mm x 250 mm, 5 μm) [57]. TBZ transformation product analysis was carried out in the samples according to previous works [58, 59] and as described in the SI.

### Modelling of enzyme-TBZ interactions

Selected ORFs which presented annotation relevant to aromatic compounds biodegradation (i.e. multi-component carbazole dioxygenase) and up-regulation in the presence of TBZ, shown in both meta-transcriptomic and meta-proteomic analysis, were computationally analyzed for possible ligand-protein interaction prognosis. Maximum common substructures between carbazole (the original substrate of the multi-component carbazole dioxygenase homologue identified as suspect catabolic enzyme in TBZ transformation) and TBZ (alternative substrate of carbazole dioxygenase, our study) were calculated with the fmcsR v1.24.0 [60] as implemented by Rcpi v1.18.1 [61]. The protein 3-dimentional structure models were calculated using SWISS-MODEL homology-based structure prediction approach [62]. Docking of TBZ was performed using Autodock Vina v1.1.2 [63] and the Autodock Tools v4.2.6 [64], while Chimera v1.11.2 [65] was used for illustration of the results.

## Results

### DNA-based SIP analysis of the bacterial consortium

The bacterial consortium was supplied with ^13^C-labelled and unlabelled TBZ as sole carbon source and the composition of the bacterial consortium was determined at four time-points along the degradation of TBZ (Figure 1A). DNA from the heavy (1.72 - 1.75 g ml^−1^) and the light fractions (1.69-1.72 g ml^−1^) of the cultures supplemented with ^13^C-labelled TBZ (Figure 1B), and total DNA from the cultures supplemented with unlabelled TBZ was subjected to amplicon sequencing. In all treatments the bacterial consortium was dominated by the same nine main OTUs belonging to α- (*Sphingomonas, Bradyrhizobiaceae, Rhizobium, Novosphingobium, Hyphomicrobium* and *Shinella*), β- (*Hydrogenophaga, Thiobacillus*) and *γ-Proteobacteria (Thiohalobacter*) (Figure 1C). However, their relative abundance varied in the heavy DNA fraction obtained from the ^13^C-TBZ supplied consortium, compared with the patterns observed in the corresponding light DNA fraction, and the consortium supplied with unlabelled TBZ (Figure 1C). Although the *Sphingomonas* OTU was dominant in all fractions and growth conditions, it entirely dominated the heavy DNA fraction of the community grown on ^13^C-labelled TBZ (>80 % relative abundance).

### Multi-omic and non-target MS analysis of TBZ transformation products

We subsequently determined in a time-series experiment (i) the meta-genome/-transcriptome/-proteome of the bacterial consortium supplemented with TBZ or succinate (offered as a non-selective C source) and (ii) the transformation products produced during the degradation of TBZ by the consortium. These results allowed us to explore possible cross-feeding associations between members of the consortium, to identify their metabolic potential and genes/enzymes with possible role in the transformation of TBZ and eventually to propose a putative transformation pathway of TBZ.

#### Metagenome analysis of the bacterial consortium

The metagenome assembly resulted in 6,742 contigs with an N50 of ~100 kb and an overall assembly length of ~98.5 Mbp (considering contigs > 500 bp; Table S1), with the devoted sequencing effort sufficiently covering the existing diversity (Nonpareil based coverage of > 99% in all samples). Binning resulted in a total of 39 MAGs with 18 of them showing a minimum of 80 % genome completeness, and 7 of them having at least 90% quality according to MiGA and therein implemented indices [41], while contamination was 1.6 % on average (± 1.4 % SD; Table S1). Amongst the assembled MAGs 11 showed mean relative abundance (in the three time points studied, 57, 96 and 120 h corresponding to 50%, 100% and 24h after complete degradation of TBZ) higher than 1% and all but one (MAG 8C, *Filimonas lacunae*, Bacteroidetes) were associated with α-, β- and *γ-Proteobacteria*, in line with the SIP-amplicon sequencing data. The six most abundant MAGs accounted for over 72% of the assembled metagenome hits and identified as γ-Proteobacterium MAG 34A (mean abundance ± standard deviation, 21.8_± 7.9%), *Sphingomonas* MAG 3X21F (17.9 ± 7.1%), *Bradyrhizobiaceae* MAG 9B (16.3 ± 9.7%), *Hydrogenophaga* MAG 23F (8.9 ± 4.2%), *Hydrogenophaga* MAG 19A (4.5 ± 6.3%) and *Hydrogenophaga* MAG 13A (3.2 ± 1.8%) (Table S1).

Translated ORF screening of the metagenome of the bacterial consortium against the AromaDeg database [43] indicated the presence of several genes scattered in the different MAGs and unbinned contigs with functional annotation associated with the aerobic degradation of monoaromatic (benzoate, genitsate, protocatechuate, salicylate, phtalate) and polyaromatic (biphenyl) hydrocarbons (Figure S1). Of particular interest was the presence of a carbazole dioxygenase (*car*), with carbazole being a structural homologue of TBZ (Figure S2), and a catechol *ortho*-cleavage pathway (*cat*) operon on MAG 3X21F contigs of *Sphingomonas*, which was identified as the main consumer of ^13^C-TBZ (Table 1). Comparative genomics between the *Sphingomonas* MAG 3X21F and other sphingomonad genomes revealed synteny between the genomic regions of *Sphingomonas* 3X21 MAG containing the *car/cat* operons and four unbinned contigs with a 50 kb region of the pCAR3 plasmid of the carbazole-degrading *Novosphingobium* sp. strain KA1 [66] (Figure 2A). Both *car* operons (in *Sphingomonas* 3X21F MAG and in *Novosphingobium* sp. KA1) were missing *carAd* encoding the ferredoxin reductase component of carbazole dioxygenase. Instead we identified in another contig of the 3X21F MAG of *Sphingomonas* a ferredoxin reductase homolog gene *fdrI* (Table 1), known to be responsible for transferring electrons from NADH^+^ to the reductase component of carbazole dioxygenase, CarAc, during carbazole transformation by *Novosphingobium* sp. KA1 [67].

**Figure 2.**
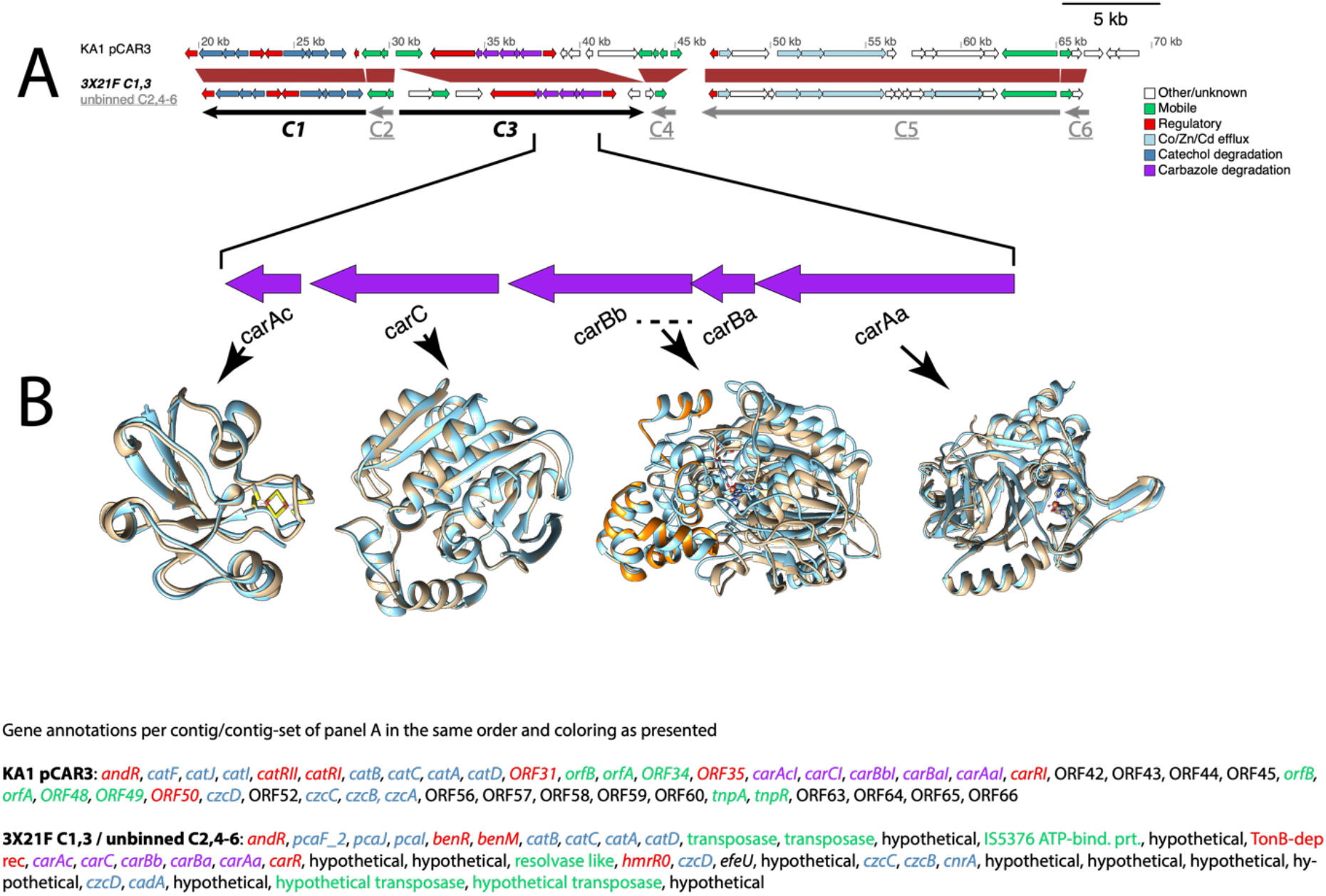
(A) Comparative analysis of a 50 kb stretch of the 255 kb long pCAR3 plasmid (upper arrow panel) carried by *Novosphingobium* sp. KA1 (NCBI accession AB270530) [66] with the *car* and *cat* operons found in the *Sphingomonas* 3X21 MAG (lower arrow panel, contigs C1 and C3),and also the unbinned contigs C2, C4, C5 and C6 of the assembled metagenome (NCBI accessions for C1-6: QFCS01001188.1, QFCS01006733.1, QFCS01001185.1, QFCS01006070.1, QFCS01006598.1, QFCS01006504.1). Red trapezoid shapes between upper and lower arrows panel denote homologous regions between the bacterial consortium metagenome and pCAR3. Annotations of all the genes in the two arrow panels are provided at the figure bottom, colored in consistency with the colors of the graphic gene representations. (B) Structural alignments of the putative carbazole operon enzymes found in *Sphingomonas* 3X21 MAG with homologous characterized protein crystal structures deposited in the protein databank (PDB; homologues are represented with the light blue crystal structures while our modeled protein structures have brown and orange colors – in the case of the CarBaBb dimer). Closest PDB accession number per gene product: 1E9M for CarAc; 1B4U for CarBaBb; 3GKQ for CarAa; 1J1I for CarC.

**Table 1.**
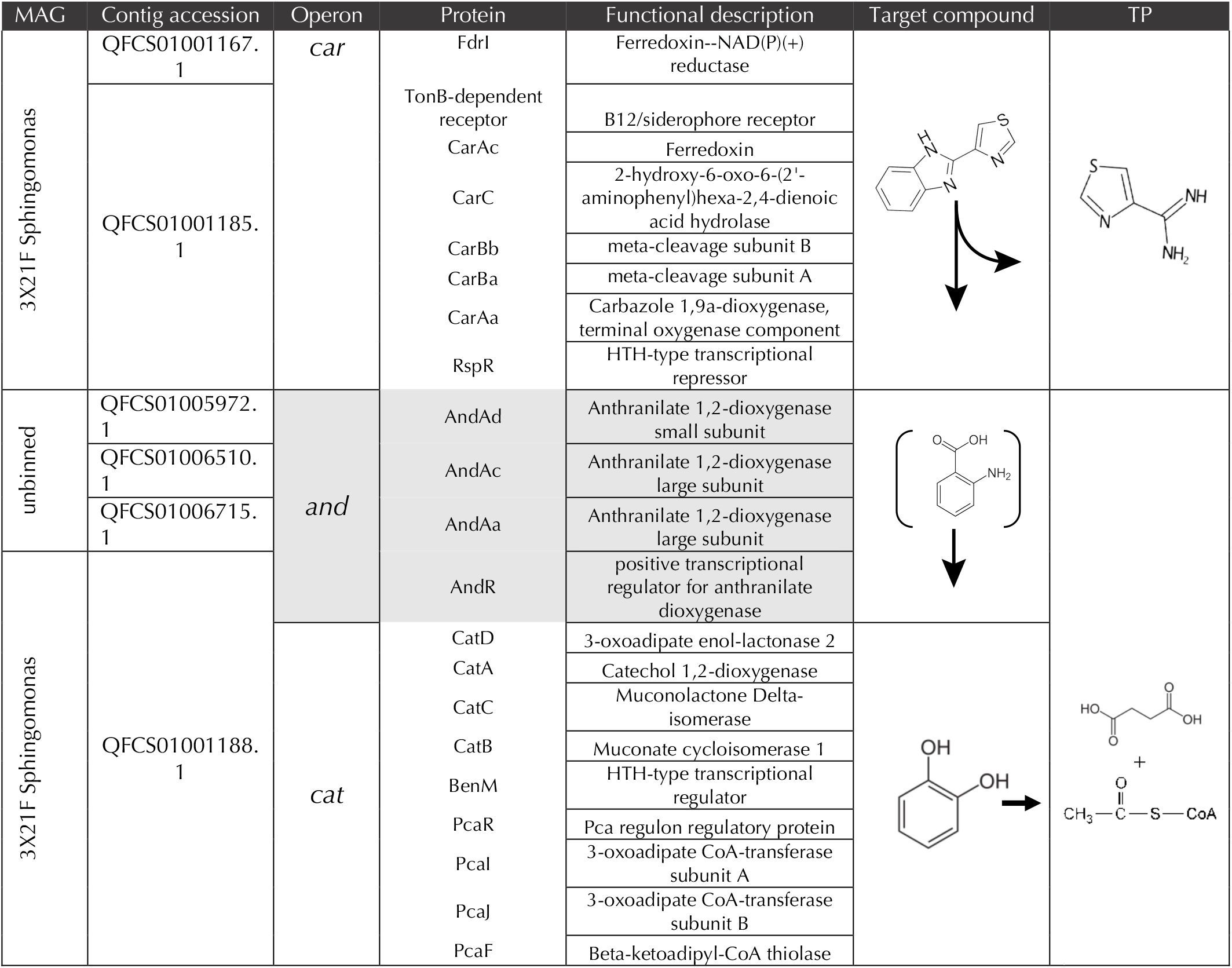
Genes and enzymes with a putative role in the transformation of thiabendazole (TBZ) as identified in the consortium metagenome. Their contig and operon residence, functional description, the target compound transformed, and the relevant transformation products derived are presented. *Car* operon encodes the enzymes responsible for the transformation of TBZ to thiazole-4-carboxamidine and catechol (directly or through the intermediate production of anthranilate) which is further transformed, through the enzymes coded by the *cat* operon, to succinate and acetyl-CoA.

#### Meta-transcriptomic/-proteomic analysis of TBZ transformation genes

Metatranscriptomic analysis showed that 21,965 genes were differentially expressed with 2,986 being up-regulated and 408 being down-regulated in the presence of TBZ compared to succinate (Figure S3). Out of the several aromatic hydrocarbon transformation annotated genes present in the consortium metagenome, the *car* and *cat* operons present in the 3X21F MAG of *Sphingomonas* were significantly up-regulated in the presence of TBZ along with the putative ferredoxin reductase *fdrI*. (Figure 3). Furthermore, regulatory elements in the immediate vicinity of the *car* and *cat* operons (e.g. TonB dependent receptor and *rspR* transcriptional repressor at the *car* operon, and *benM/pcaR* at the *cat* operon) were also up-regulated in the presence of TBZ. We also observed a significant up-regulation of the different components of anthranilate dioxygenases (*andAcAdAa*) and their transcriptional regulator (*andR*) under TBZ supplementation (Figure 3). Apart from *andR* which was located upstream of the *cat* operon, all other anthranilate dioxygenase components were scattered in unbinned contigs (locus tags 04040, 06397, 07193; Table 1).

**Figure 3.**
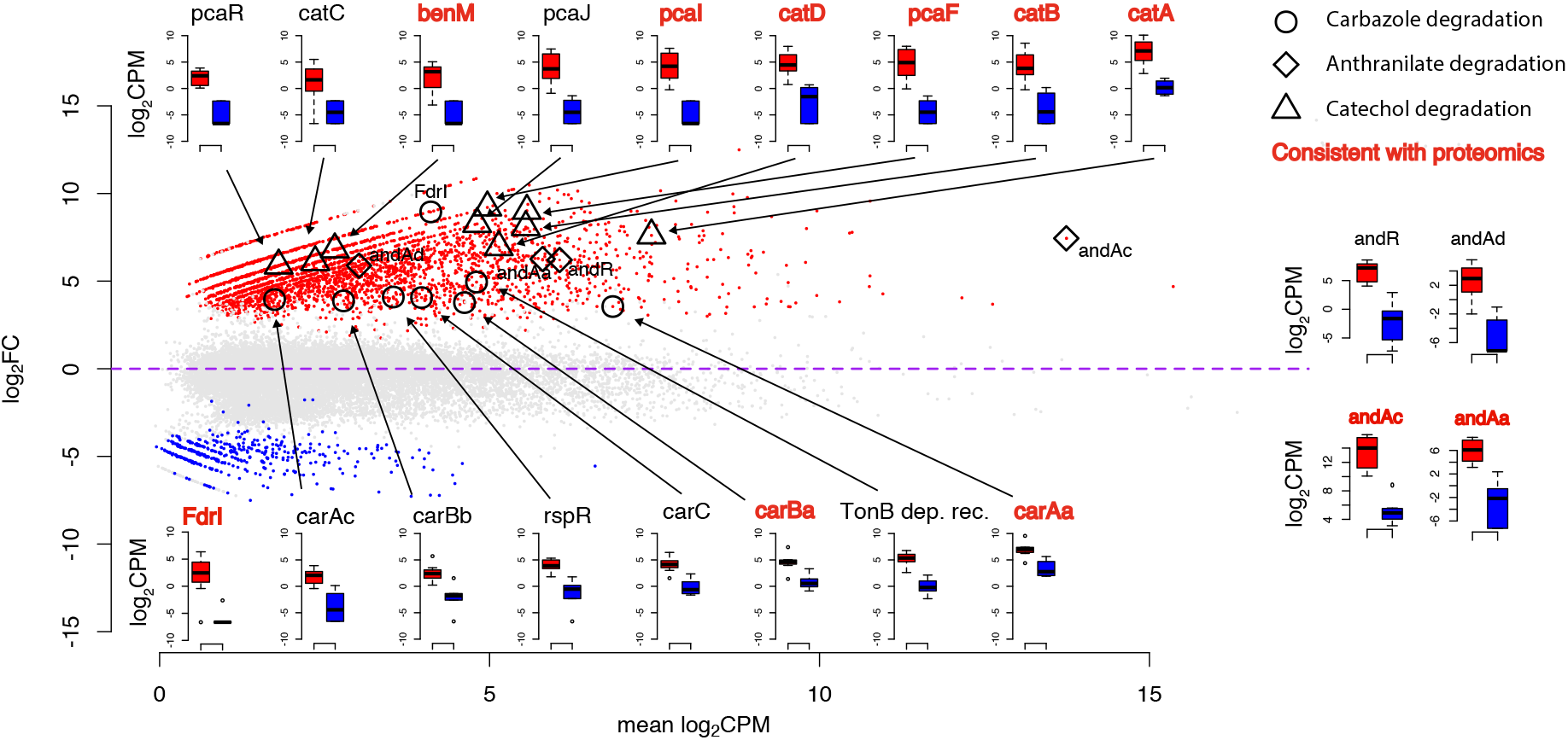
Differential expression profile of genes and enzymes of the bacterial consortium grown with thiabendazole (TBZ) or succinate. Data are presented as log2(fold change) *vs* the means of the log2(copies per million reads - CPM) (MA) plot of the treatment-related gene differential expression according to RNA. The *car*, *cat* and *and* loci associated points are depicted with open circles, triangles and diamonds on the plots respectively. The expression boxplots in log2CPM values for genes with a putative role in the transformation of TBZ are also provided with arrows and/or symbols connecting them to the points of the MA plot. Genes/enzymes which showed consistent differentially expressed patters at both metatranscriptomics and metaproteomics analysis are depicted with bold red letters according to the key.

Corresponding meta-proteomic analysis of the bacterial consortium showed 2602 proteins to be differentially expressed in the two feeding conditions, 423 proteins being up-regulated and 652 down-regulated in the presence of TBZ (Figure S3). When focused on proteins with putative role in the transformation of TBZ, we noticed that the translated products of several of the genes of the *car* (FdrI, CarAaBa) and *cat* (CatABD, PcaIF) operons showed (consistent to the meta-transcriptomic data) significantly up-regulated profiles in the presence of TBZ (Figure 3). A consistent up-regulated profile in the presence of TBZ was also evident for anthranilate dioxygenase components AndAaAc (Figure 3).

Modeling and structural comparisons of the CarAaAcBaBbC components of the carbazole dioxygenase locus (Figure 2B) and of FdrI (Figure S4) found in the *Sphingomonas* 3X21F MAG with the corresponding components of the homologous characterized carbazole dioxygenase components of *Novosphingobium* sp KA1, showed nearly identical three-dimensional conformations. CarAa showed 77% identities and 87% positives over the complete translated ORF length with the *Novosphingobium* sp. KA1 putative homologue and 30 identical out of the 32 amino acids around the active site pocket (Figure S5). The predicted CarAa active site pocket had similar affinity to carbazole (ΔG = −7.5 kcal mol^−1^) compared to previously characterized CarAa enzymes (corresponding ΔGs of −8.4 and −7.4 kcal mol^−1^ for *Janthinobacterium* and *Novosphingobium* sp. KA1 respectively), and a slightly lower affinity to TBZ (corresponding ΔG values of −6.8 kcal mol^−1^ compared to ΔG values of −7 and −6.6 kcal mol^−1^ for *Janthinobacterium* and *Novosphingobium* sp. KA1 respectively) (Figure S6).

### Transcriptional profile associations between consortium members

We further investigated potential interdependencies and associations at functional level between the different members of the consortium. We performed a differentially-expressed-gene network analysis based on the metatranscriptomic data. We identified two dominant network substructures with connections between genes of *Sphingomonas* MAG 3X21F and the MAGs classified as *Hydrogenophaga* 23F, *Bradyrhizobiaceae* 9B, and unbinned contigs. Correlation analysis between the SEED functional gene categories of the genes comprising the network substructures, revealed a coincident expression of the genes contained in the putative *car* and *cat* operons of *Sphingomonas* MAG 3X21F and genes encoding cobalamin biosynthesis and its transmembrane transportation in the *Hydrogenophaga* MAG 23F (Figure 4A). TonB-dependent transporter genes which are responsive to siderophores, colisin and cobalamin [68], signaling genes with homology to the *fixLJ* two component systems (9 ORFs) and secondary messengers like cyclic AMP were all up-regulated in the presence of TBZ only in the *Sphingomonas* MAG 3X21F, although only cAMPs expression correlated with other MAGs (Figure 4A).

**Figure 4.**
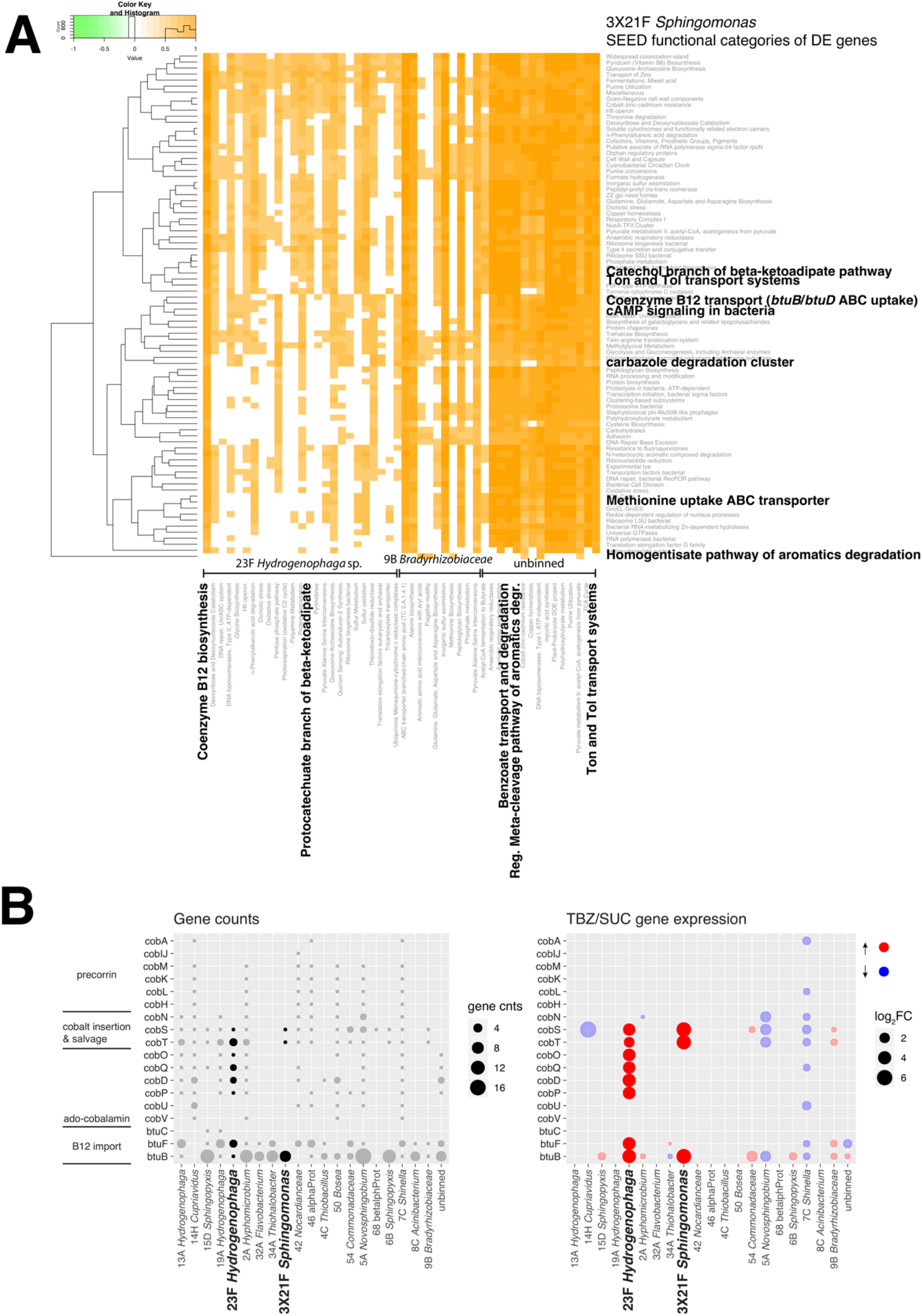
(A) Heatmap of the correlations (Spearman ρ ≥ 0.05, and Bejamini-Hochberg adjusted P-value cutoff of ≤ 0.01) of the differentially expressed genes (Bejamini-Hochberg adjusted P-value cutoff of ≤ 0.01) of the two dominant network substructures including *Sphingomonas* 3X21F MAG genes, against the differentially expressed of other MAGs or genes on unbinned contigs of the same network groups. (B) Bubble plot showing the presence of cobalamin biosynthesis and transportation gene counts in the different MAGs of the bacterial consortium metagenome (left) and the expression profile (up- or down-regulated) of significantly differentially expressed genes when supplemented with TBZ or succinate (SCU) (log2(TBZ/SUC); right). Per-panel provided keys explain significance of the plotted colors and bubble sizes.

We further investigated the completeness and the expression profile of the B12 biosynthesis and transportation systems encoded in the different MAGs (Figure 4B). Most major MAGs including *Sphingomonas* 3X21F contained several copies of the *btuB* and *btuF* genes encoding modules of the cobalamin transmembrane translocation system [73]. Regarding cobalamin biosynthesis *Hydrogenophaga* MAG23F carried copies of several of the genes necessary for the biosynthesis of the co-factor, while near complete B12 biosynthesis pathways were noted on other MAGs like *Hyphomicrobium* MAG 2A, *Shinella* MAG 7C and *Novosphingobium* MAG 5A (Figure 4B). On the other hand, *Sphingomonas* MAG 3X21F was amongst the poorest of the MAGs in genes associated with cobalamin biosynthesis. When the differential expression of the genes associated with the biosynthesis and transportation of cobalamin in the different MAGs was recorded, we observed an up-regulation of the full array of the relevant genes in *Hydrogenophaga* MAG23F with concurrent up-regulation of genes *btuB* and *cobST* associated with transmembrane transportation and salvage of cobalamin respectively in *Sphingomonas* MAG 3X21F [69–71] (Figure 4B). In contrast we observed down-regulation of the corresponding *cob* genes in other MAGs (*Hyphomicrobium* 2A, *Novosphingobium5A* and *Shinella* 7C) with a populated biosynthetic pathway of cobalamin.

Assessment of keystoneness indices [54] based on the metatranscriptomic data network analysis further demonstrated the central role of the *Sphingomonas* MAG 3X21F among other MAGs (Figure S7). The genes of MAG 3X21F showed the highest degree, indirect degree (significance group a), and transitivity (highest significance groups; Figure S7), a relatively high closeness centrality (ranked 4^th^ out of 9 significance groups) and the lowest betweenness centrality values (8 out of 8 significance groups).

### Non-target MS analysis of TBZ transformation products

In the same time series experiment (experiment 2) we determined, via non-target MS analysis, the formation of potential transformation products of TBZ. The rapid degradation of TBZ was accompanied by the concurrent formation of a single major metabolic product with an MS spectrum of 127 m/z [14, 57–59] identified as 1,3-thiazole-4-carboxamidine (Figure 5A). This product was not further transformed by the bacterial consortium. We also detected two other minor and transient transformation products in the bacterial culture identified as 5-OH-thiabendazole, with an MS spectrum of 218 (Figure 5B), and thiazole carboxamide, with an MS spectrum of 128 m/z (Figure 5C) (MS spectra given in our previous studies with TBZ [14, 57–59]).

**Figure 5.**
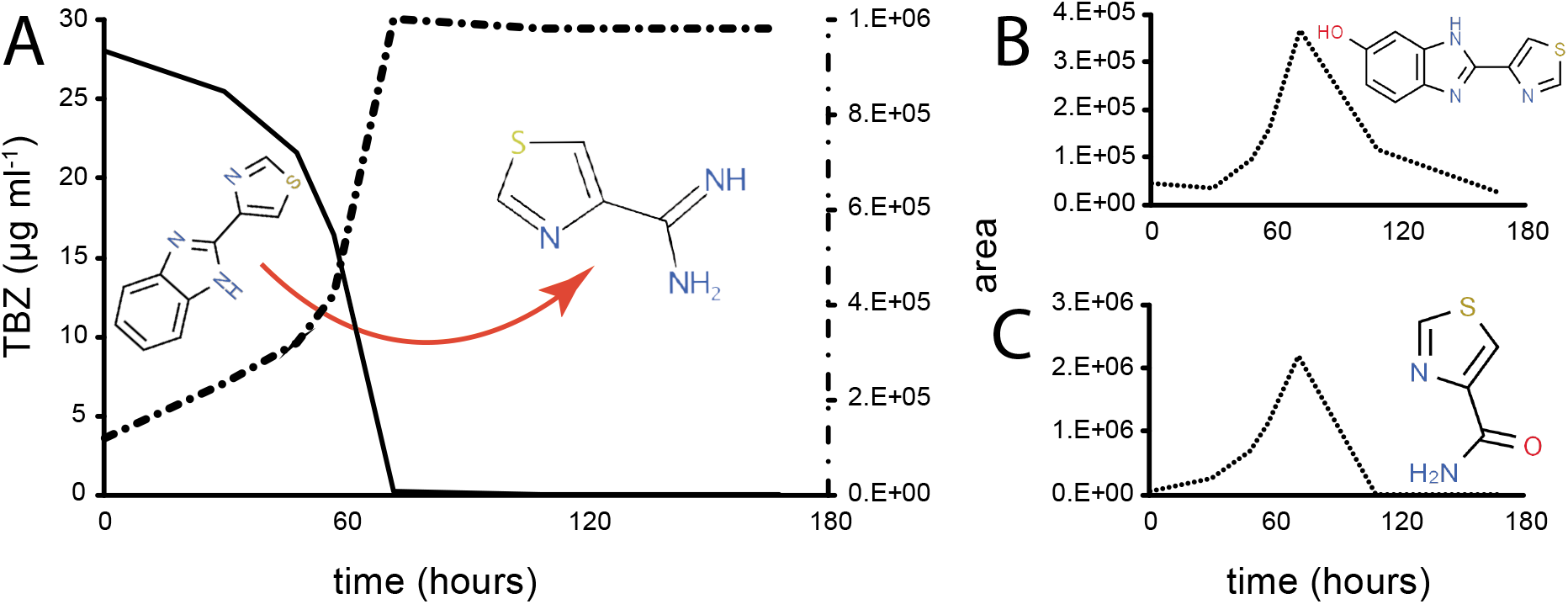
(A) The degradation of thiabendazole (TBZ) by the bacterial consortium (solid line, μg ml^−1^) and the formation (MS peak area) of the persistent transformation product 1,3-thiazole-4-carboxamidine (dash-dotted line); (B) and (C) The formation and decay of the MS peak areas of the transient transformation products 5-OH-thiabendazole and thiazole-4-carboxamide respectively.

## Discussion

Bacterial consortia often encompass complex metabolic and nutritional interactions which support effective pollutant degradation and their coherence. Working previously with a TBZ-degrading consortium we showed, via SIP-DGGE analysis at a single time point (taken upon completion of TBZ degradation) that a *Sphingomonas* sp. was the main TBZ degrader with the contribution of a *Hydrogenophaga* being suggested [14]. Here, by employing a time-series SIP-based amplicon sequencing approach we verified that *Sphingomonas* sp. is the sole member of the consortium involved in the transformation of TBZ, and no cross-feeding events of transformation products were observed.

We also assembled the metagenome of the bacterial consortium aiming to unravel the metabolic potential of its individual members. The consortium metagenome was composed of 18 major MAGs (with at least 80 % completeness); of those six were dominant, when grown on TBZ, with a rich arsenal of genes encoding the transformation of aromatic organic pollutants. Focusing on MAG 3X21F of *Sphingomonas* as the TBZ degrader of the consortium, we found genes with hits in the benzoate, biphenyl, extradiol, gentisate, LigB, protocatechuate and phthalate dioxygenase (super) families [43]. Most interestingly, we noticed two DNA stretches approximating 20-kb, encompassing a *car* and a *cat* operon, both being significantly up-regulated during TBZ degradation according to meta-transcriptomic and meta-proteomic analysis, suggesting their involvement in the transformation of TBZ. Carbazole is an *N*-heterocyclic aromatic hydrocarbon, structurally similar to TBZ (Figure S2), found in fossil fuels and coal/wood combustion products, and it is used as a chemical feedstock for the production of dyes, reagents, explosives, insecticides, lubricants [72]. The carbazole skeleton is also present in natural alkaloids produced by the roots of several plants [73]. The high anthropogenic and biogenic exposure of several environments to carbazole skeleton explains the evolution of relevant catabolic degrading systems and their widespread occurrence amongst bacteria isolated from activated sludge and soil [74–76]. The *car* locus codes enzymes for the transformation of carbazole (and several other aromatic pollutants) to catechol through the intermediate production of anthranilate [67]. The full transformation pathway includes the electron transfer from NAD(P)H through a ferredoxin reductase (FdrI/FdrII in *Novosphingobium* sp. KA1 [67] and CarAd in *Pseudomonas resinovorans* CA10 [77]) and a ferredoxin (CarAc) to the terminal oxygenase component (CarAa) which angularly dioxygenates carbazole. Following, a spontaneous cleavage takes place to form 2’-aminobiphenyl-2,3-diol, which is further cleaved by CarBaBb to a *meta*-cleavage product (2-hydroxy-6-oxo-6-(2’-aminophenyl)-hexa-2,4-dienoic acid), further hydrolyzed to anthranilate and 2-hydroxypenta-2,4-dienoic acid by CarC. The former is transformed to catechol by anthranilate dioxygenase AndAaAbAd and finally to succinate and acetyl Co-A via the catechol *ortho*-cleavage pathway. The *car* operon found in *Sphingomonas* MAG 3X21F comprised *carAaAcBaBbC*, lacking *carAd*. This is a common feature of other carbazole-degrading sphingomonads like *Sphingomonas* strain XLDN2-5 [78] and *Novosphingobium* sp., GTIN11 [79] but not of *Pseudomonas* CA10 [80] and *Nocardioides aromaticivorans* IC177 [81] where *carAd* is part of the *car* operon. In the absence of *carAd* in its *car* operon, *Novosphingobium*sp. KA1 plasmid pCAR3 contains genes coding for two ferredoxin reductases, FdrI/FdrII, located away from the two *car* operons of *Novosphingobium* sp. KA1and acting as CarAd substitutes [67, 82]. Screening of *Sphingomonas* MAG 3X21F contigs lead to the identification of an FdrI homolog, which was highly up-regulated under TBZ at transcript and protein levels. The *fdrI* ORF found on the MAG of the main degrader, showed differential expression in the presence of TBZ, contradicting the previously reported constitutive expression in *Novosphingobium* sp. KA1 plasmid pCAR3 [82]; the difference in the expression pattern may be related to the range of the enzyme-dependent reactions in each tested case. Previous tests have also shown a broad range of activity compensation of FdrI/FdrII by spinach and *Escherichia coli* crude extract ferredoxin reductases by 96% and 4.5% respectively [77]. These results and the observed genetic organization of the relevant genes might suggest that *fdrI* is shared amongst several catabolic pathways, as previously proposed to be an evolutionary strategy of sphingomonads to maximize their catabolic potential and minimize their energetic burden [82, 83]. Alternatively, the pathway could be still under evolution [82]. The structural similarity of TBZ with carbazole, the remarkably relaxed specificity of carbazole dioxygenase which could transform a very wide range of polyaromatic pollutants (i.e., dibenzofuran, dibenzo-p-dioxin, biphenyl, napthalene, dibenzothiophene, diphenylamine) [67, 84] and its consistent up-regulation while growing with TBZ, suggest that carbazole dioxygenase is responsible for the initial step of the transformation of TBZ by the bacterial consortium. Further support of this is provided by the almost equivalent affinity of the CarAa found in 3X21F MAG of *Sphingomonas* to carbazole and TBZ indicated by *in silico* docking tests.

Comparative analysis between the whole consortium metagenome and carbazole-degrading bacterial genomes showed high synteny of the *car/cat* operons with a 50 kb stretch of the pCAR3 plasmid encoding the full pathway for the transformation of carbazole in *Novosphingobium* sp. KA1 [66]. Both catabolic operons in MAG 3X21F and in *Novosphingobium* sp. KA1 were flanked by transposable elements suggesting that the *car/cat* genomic region of MAG 3X21F of *Sphingomonas* is a patchy construction acquired most probably by other sphingomonads through horizontal gene transfer. Sphingomonads are considered as “artists of biodegradation” due to their versatility in the degradation of organic pollutants [85–87] stemming from their remarkable capacity to exchange whole plasmids or mobile genetic elements to expand their catabolic repertoire [88, 89].

The *andAaAbAd* genes, although upregulated under the TBZ treatment, were not organized in the vicinity of the *car/cat* operons in MAG 3X21F, compared to the pCAR3 plasmid of *Novosphingobium* sp. strain KA1 where *and* genes were located closely upstream of the *cat* operon and in close proximity to the *car* operon [66]. Instead, in our case they were scattered in MAG-unassigned contigs implying that anthranilate is not produced during transformation of TBZ, in line with its lack of detection in the culture during TBZ degradation. However, it cannot be excluded that its presence in unbinned contigs might be associated with limitations of the metagenome assembly and binning methods with respect to the available sequencing technologies. For instance, the existence of several homologs of anthranilate dioxygenase throughout the consortium members (considering its role in the degradation of tryptophan and other aromatic compounds [84]) and its common localization in transposable elements [66, 78], may have resulted in population level repetitive regions which are usually recalcitrant to contiguousness in assembly, and also to coverage and sequence structure based binning [90]. In contrast, we found a complete putative *cat* operon composed of *catABCD* and *pcaIJF*, binned in MAG 3X21F of

*Sphingomonas* which was consistently upregulated, at both transcription and protein level, suggesting its operation in the transformation of catechol produced by the phenyl moiety of the benzimidazole ring of TBZ to tricarboxylic acid (TCA) cycle intermediates.

Considering all our meta-omic analysis, non-target MS analysis, current and previous isotopic (^13^C and ^14^C) studies [14] we propose a transformation pathway of TBZ fully accomplished by *Sphingomonas* 3X21F (Table 1): TBZ is initially cleaved at the imidazole moiety of the benzimidazole ring by carbazole dioxygenase CarAaAcFdr; this produces an unknown dioxygenated transient intermediate which is *meta*-cleaved by CarBaBb and hydrolyzed by CarC to 1,3-thiazole-4-carboxamidine as a dead-end transformation product (in line with previous findings [14] and our non-target MS analysis), and catechol (directly or through the intermediate production of anthranilate). This is further transformed by *Sphingomonas* 3X21F (in line with the assimilation of the ^13^C-phenyl moiety of TBZ in the SIP analysis and previous radiorespirometric assays [15]) probably to the *ortho*-cleavage pathway terminal products acetyl-CoA and succinate, entering the TCA cycle.

We further looked for nutritional interdependencies between consortium members, beyond the transformation of TBZ, that might contribute to the consortium coherence. RNA-based network analysis revealed that certain functional features of *Sphingomonas* MAG 3X21F were highly correlated with *Hydrogenophaga* MAGs 23F, *Bradyrhizobiaceae* MAG 9B and unbinned contigs. Amongst them we observed a prominent positive correlative expression between cobalamin biosynthesis in *Hydrogenophaga* MAGs 23F and the catabolism of carbazole/catechol or the transportation system of cobalamin in *Sphingomonas* MAG 3X21F. Although several members of the consortium appear to possess a well populated cobalamin biosynthetic pathway, transcriptomic analysis revealed that only *Hydrogenophaga* MAG 23F mobilizes its cobalamin biosynthetic pathway (CobSTOQDP) and trans-membrane transportation, (BtuBF) in the presence to TBZ (Figure 6B). At the same time *Sphingomonas* MAG 3X21F, which is probably not capable of *de novo* biosynthesis of cobalamin, has activated its cobalamin transmembrane transportation system (BtuBF) and the *CobST* genes participating in the salvage pathway of cobamides [69, 70]. Since there are no known altruistic B12-producing bacteria [22], our findings indicate a nutritional inter-dependency between *Sphingomonas* 3X21F and *Hydrogenophaga* 23F, complementing the probable B12 auxotrophy of the former with the B12 prototrophy of the latter. Several studies have stressed the key role of cobalamin in shaping microbial communities [91], based on its key role as a co-factor in the reductive dehalogenation of organic pollutants [92] and other key metabolic reactions, and the evolutionary loss of the energetically costly B12 biosynthesis by most bacteria [23, 93, 94]. The missing cobalamin biosynthetic genes from the *Hydrogenophaga* MAG 23F could be the result of possible sequence divergence from characterized database genes or a low coverage of these regions by our sequencing effort.

An interesting point is the specificity of the correlations between *Sphingomonas* MAG 3X21F and the *Hydrogenophaga* MAG 23F, given the presence of other *Hydrogenophaga* MAGs in the bacterial consortium metagenome which did possess cobalamin biosynthetic genes but showed no transcriptional correlation with the *Sphingomonas* MAG 3X21F. Hence, we looked for the activation of potential signaling mechanisms involving autoinducers or secondary messengers which might trigger this specific interaction. In the presence of TBZ we observed up-regulation of several *luxRI*-like two-component sensory-histidine-kinase/regulatory genes, annotated as *fixLJ* [95, 96] at the *Sphingomonas* MAG 3X21F, which did not appear to correlate with other members of the consortium. The *fixLJ* two component system is considered to sense oxygen in nitrogen fixing bacteria, yet, a broader sensory role of this two-component system was postulated in other bacteria (i.e. in the virulence of *Bulkhorderia dolosa*) [97, 98]. Cyclic AMPs, on the other hand, are involved mainly in intracellular signaling and were recently shown to participate in generic extracellular signaling, particularly during stress conditions [99, 100]. These two signaling modes were upregulated under the TBZ treatment and participated in the correlation network between the *Sphingomonas* MAG 3X21F and MAG 23F of *Hydrogenophaga*. These traits and our results render them suitable candidate systems mediating an interaction between the strains represented by the corresponding MAGs, which may have resulted in the MAG 23F derived cobalamin support of *Sphingomonas* (MAG 3X21F). A possible alternative mode for triggering cobalamin biosynthesis in *Hydrogenophaga* 23F could be associated with stress imposed by the presence of TBZ known to exert toxic effects on prokaryotes [101–103]. Besides *Sphingomonas*

MAG 3X21F, showing up-regulation of several stress-related genes when grown on TBZ (39 genes), a pattern often observed in pollutant-degrading bacteria during exposure to the target pollutant [104–106], *Hydrogenophaga* 23F was the only other MAG that mobilized a stress response, involving the up-regulation of a general stress response gene and a glutathione-S-transferase gene (locus tag: Bin_23F_Hydrogenophaga_01167/03148). Oxidative stress elicitors have been shown to stimulate cobalamin production in bacteria [107]. Yet, this would also fail to explain on its own the specificity of the interaction, unless there are pronounced metabolic and functional differences among *Hydrogenophaga* consortium members, previously reported for *Hydrogenophaga* isolates [108], which enable this specialized interaction to occur.

## Conclusions

We previously demonstrated the efficient degradation of the fungicide TBZ by a bacterial consortium and showed first evidence for the role of a *Sphingomonas* in TBZ degradation [14]. Here we provide unequivocal evidence that *Sphingomonas* is degrading TBZ without the involvement of any other member of the consortium. To achieve this, it employs a carbazole dioxygenase and a catechol *ortho*-cleavage operon, co-localized in a composite transposon most probably derived through horizontal gene transfer by other sphingomonads upon selection pressure imposed by TBZ contamination. In contrast to the lack of metabolic cross-feeding among consortium members in the transformation of TBZ, we observed a strong metabolic interdependency of the B12 auxotroph *Sphingomonas*, based on its assembled genome, with a *Hydrogenophaga* 23F which activates its cobalamin biosynthesis in response to TBZ degradation. Based on our results we propose a model, schematically presented in Figure 6, that describes the function of the TBZ-degrading consortium in line with the “black queen hypothesis” [109]: *Sphingomonas* 3X21F, is taking over the degradation of TBZ, relieving consortium members by a prokaryotic toxicant like TBZ [101–103]. In exchange for this, *Hydrogenophaga* 23F provides cobalamin to the auxotrophic *Sphingomonas* 3X21F ensuring the survival of the “black queen” of the consortium.

**Figure 6.**
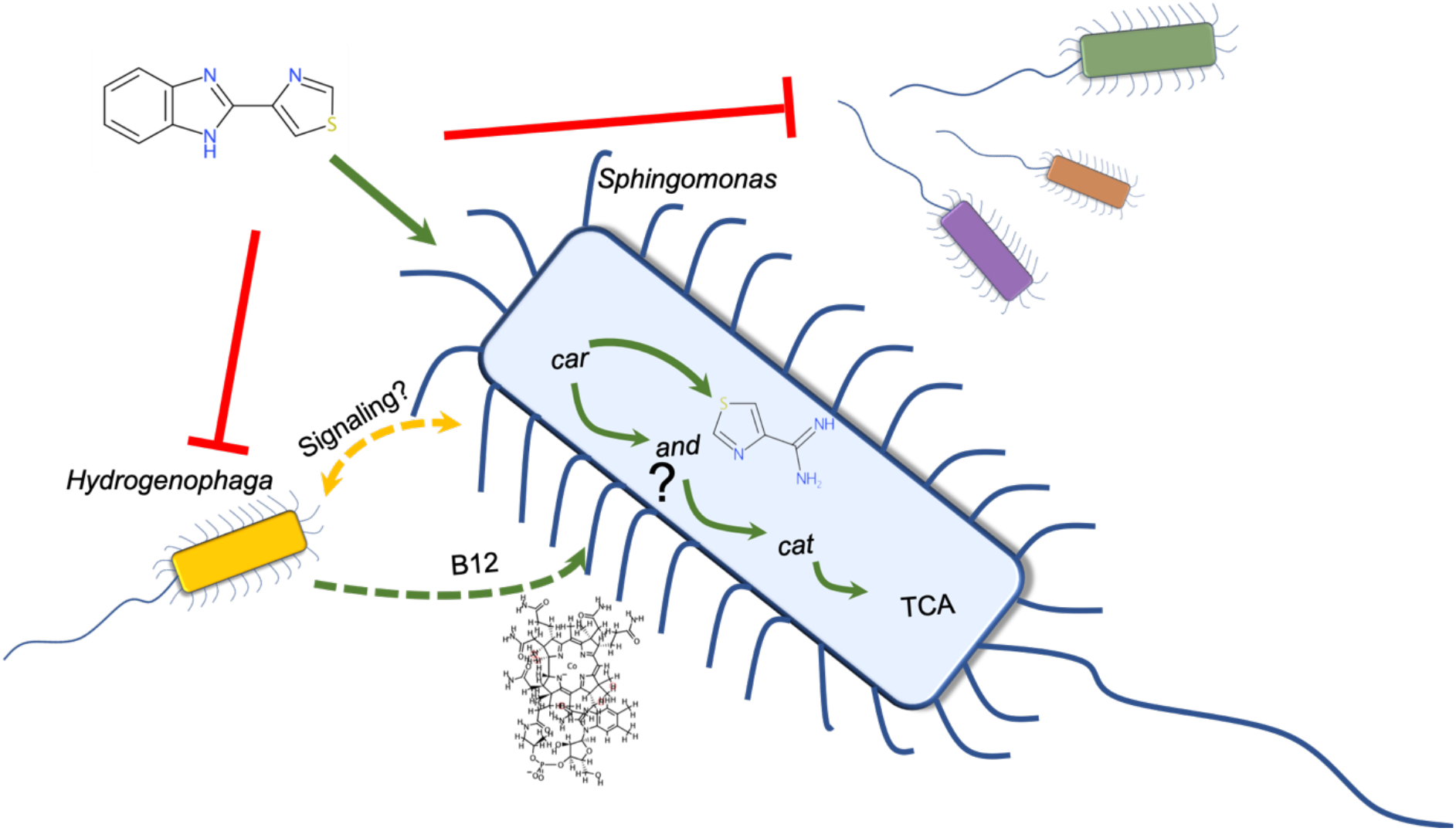
Proposed model describing the interactions between the members of the bacterial consortium driving the degradation of thiabendazole (TBZ). Green arrows indicate putative nutrient direction flows; yellow double arrows indicate signaling interactions; red blocked lines indicate possible inhibitory/toxic effects. car: carbazole-dioxygenase operon; and: anthranilate dioxygenase gene set; cat: catechol *ortho*-cleavage operon; TCA: tricarboxylic acid cycle.

## Ethics approval and consent to participate

Not applicable.

## Consent for publication

All the authors have seen and agree with the contents of this manuscript.

## Availability of data and material

All sequence data are available at National Center for Biotechnology Information (NCBI) sequence read archive (SRA) under the bioproject accession number PRJNA466717 with: the metagenome assembly sequencing data accessible under the numbers SRR7135606-12; the RNA sequencing data and complete metadata are accessible through the NCBI Gene Expression Omnibus (GEO) under the accession GSE134575 and SRA through accessions SRR9719617-34; the 16S rRNA gene amplicon sequencing data accessible under the numbers SRR9699065-175. The metagenome assembly contig set is publicly available at the NCBI Genbank database with the accession number GCA_006513095.1.

## Competing interests

The authors declare no competing interests.

## Funding

Dr Vasileiadis is funded by the MSCA-IF-H2020 project EMIGRATE (Grant Agreement No. 749463), Dr. C. Perruchon and Dr D.G. Karpouzas are supported by OMIC-ENGINE (MIS 5002636) funded by the Operational Program “Competitiveness, Entrepreneurship and Innovation” (NSRF 2014-2020) and co-financed by Greece and the European Union (European Regional Development Fund). Protein mass spectrometry was done at the Centre for Chemical Microscopy (ProVIS) at the Helmholtz Centre for Environmental Research - UFZ, which is supported by European regional development funds (EFRE—Europe Funds Saxony) and the Helmholtz Association.

## Author’s contributions

KGD has conceived, designed and supervised all experiments along with the supervision of the drafting of the manuscript. PC participated in the experimental design, and the performance and supervision of the experiments. SB and AL designed the shotgun proteomics experiment and generated the data. SN and CA designed the SIP experiment and supervised its performance.PBP and AA designed the non-target MS analysis and generated the associated data. TM has participated in the experimental design and along with AL, CA and AA have provided feedback on the manuscript draft. VS has drafted the manuscript, performed all experiments except for the non-target MS analysis and the shotgun proteome analytical parts, performed the amplicon library prep for multiplex sequencing, the meta-genomics/transcriptomics/proteomics sample prep for sequencing or analysis, the TBZ transformation products library preparation, the bioinformatics, the statistics and protein-ligand docking modeling, and participated in the experimental design.

## Acknowledgements

The authors are thankful to Dr. Georgios E. Papadopoulos for his stewardship on the protein-ligand docking analysis.

## References

1. D’Aquino S, Palma A, Angioni A, Schirra M. Residue Levels and Efficacy of Fludioxonil and Thiabendazole in Controlling Postharvest Green Mold Decay in Citrus Fruit When Applied in Combination with Sodium Bicarbonate. J Agric Food Chem. 2013; 61:296–306.

2. Panic G, Duthaler U, Speich B, Keiser J. Repurposing drugs for the treatment and control of helminth infections. International Journal for Parasitology: Drugs and Drug Resistance. 2014; 4:185–200.

3. Özkay Y, Tunalı Y, Karaca H, Işıkdağ İ. Antimicrobial activity of a new series of benzimidazole derivatives. Archives of Pharmacal Research. 2011; 34:1427.

4. Zhou Y, Xu J, Zhu Y, Duan Y, Zhou M. Mechanism of action of the benzimidazole fungicide on *Fusarium graminearum:* Interfering with polymerization of monomeric tubulin but not polymerized microtubule. Phytopathology. 2016; 106:807–13.

5. Abongwa M, Martin RJ, Robertson AP. A brief review on the mode of action of antinematodal drugs. Acta Veterinaria. 2017; 67:137–52.

6. Ccanccapa A, Masiá A, Andreu V, Picó Y. Spatio-temporal patterns of pesticide residues in the Turia and Júcar Rivers (Spain). Sci Total Environ. 2016; 540:200–10.

7. Masiá A, Campo J, Vázquez-Roig P, Blasco C, Picó Y. Screening of currently used pesticides in water, sediments and biota of the Guadalquivir River Basin (Spain). J Hazard Mater. 2013; 263:95–104.

8. European Food Safety Authority. Conclusion on the peer review of the pesticide risk assessment of the active substance thiabendazole. EFSA J. 2014; 12:3880.

9. Papadopoulou ES, Tsachidou B, Sułowicz S, Menkissoglu-Spiroudi U, Karpouzas DG. Land spreading of wastewaters from the fruit-packaging industry and potential effects on soil microbes: effects of the antioxidant ethoxyquin and its metabolites on ammonia oxidizers. Appl Environ Microbiol. 2016; 82:747–55.

10. US EPA. Registration eligibility decision (RED): Thiabendazole. 2002; EPA-738-F-02-002, US Environmental Protection Agency

11. European Commission. Draft renewal assessment report prepared according to the Commission Regulation (EU) N.1141/2010, Second programme for the renewal of the inclusion of the following active substance under Regulation (EC) 1107/2009,Thiabendazole, Vol. 1, Report and proposed decision. 2013;

12. Campo J, Masiá A, Blasco C, Pico Y. Occurrence and removal efficiency of pesticides in sewage treatment plants of four Mediterranean river basins. J Hazard Mater. 2013; 263:146–57.

13. Walters E, McClellan K, Halden RU. Occurrence and loss over three years of 72 pharmaceuticals and personal care products from biosolids–soil mixtures in outdoor mesocosms. Water Res. 2010; 44:6011–20.

14. Perruchon C, Chatzinotas A, Omirou M, Vasileiadis S, Menkissoglou-Spiroudi U, Karpouzas DG. Isolation of a bacterial consortium able to degrade the fungicide thiabendazole: the key role of a *Sphingomonas* phylotype. Appl Microbiol Biotechnol. 2017; 101:3881–93.

15. Perruchon C, Pantoleon A, Veroutis D, Gallego-Blanco S, Martin-Laurent F, Liadaki K, et al. Characterization of the biodegradation, bioremediation and detoxification capacity of a bacterial consortium able to degrade the fungicide thiabendazole. Biodegradation. 2017; 28:383–94.

16. Breugelmans P, Barken KB, Tolker-Nielsen T, Hofkens J, Dejonghe W, Springael D. Architecture and spatial organization in a triple-species bacterial biofilm synergistically degrading the phenylurea herbicide linuron. FEMS Microbiol Ecol. 2008; 64:271–82.

17. Kim H, Kim D-U, Lee H, Yun J, Ka J-O. Syntrophic biodegradation of propoxur by Pseudaminobacter sp. SP1a and Nocardioides sp. SP1b isolated from agricultural soil. Int Biodeterior Biodegrad. 2017; 118:1–9.

18. Men Y, Feil H, VerBerkmoes NC, Shah MB, Johnson DR, Lee PKH, et al. Sustainable syntrophic growth of *Dehalococcoides ethenogenes* strain 195 with *Desulfovibrio vulgaris* Hildenborough and *Methanobacterium congolense:* global transcriptomic and proteomic analyses. ISME J. 2011; 6:410.

19. Hug LA, Beiko RG, Rowe AR, Richardson RE, Edwards EA. Comparative metagenomics of three Dehalococcoides-containing enrichment cultures: the role of the non-dechlorinating community. BMC Genomics. 2012; 13:327.

20. Xu X, Zarecki R, Medina S, Ofaim S, Liu X, Chen C, et al. Modeling microbial communities from atrazine contaminated soils promotes the development of biostimulation solutions. ISME J. 2019; 13:494–508.

21. Mee MT, Collins JJ, Church GM, Wang HH. Syntrophic exchange in synthetic microbial communities. Proc Natl Acad Sci. 2014; 111:E2149–E56.

22. Shelton AN, Seth EC, Mok KC, Han AW, Jackson SN, Haft DR, et al. Uneven distribution of cobamide biosynthesis and dependence in bacteria predicted by comparative genomics. ISME J. 2019; 13:789–804.

23. Zengler K, Zaramela LS. The social network of microorganisms — how auxotrophies shape complex communities. Nat Rev Microbiol. 2018; 16:383–90.

24. Mee MT, Wang HH. Engineering ecosystems and synthetic ecologies. Molecular BioSystems. 2012; 8:2470–83.

25. De Vrieze J, Boon N, Verstraete W. Taking the technical microbiome into the next decade. Environ Microbiol. 2018; 20:1991–2000.

26. Neufeld JD, Vohra J, Dumont MG, Lueders T, Manefield M, Friedrich MW, et al. DNA stable-isotope probing. Nat Protocols. 2007; 2:860–6.

27. Vasileiadis S, Puglisi E, Trevisan M, Scheckel KG, Langdon KA, McLaughlin MJ, et al. Changes in soil bacterial communities and diversity in response to long-term silver exposure. FEMS Microbiol Ecol. 2015; 91:fiv114.

28. Vasileiadis S, Puglisi E, Papadopoulou ES, Pertile G, Suciu N, Pappolla RA, et al. Blame it on the metabolite: 3,5-dichloraniline rather than the parent compound is responsible for decreasing diversity and function of soil microorganisms. Appl Environ Microbiol. 2018; 84:e01536–18.

29. Parada AE, Needham DM, Fuhrman JA. Every base matters: assessing small subunit rRNA primers for marine microbiomes with mock communities, time series and global field samples. Environ Microbiol. 2016; 18:1403–14.

30. Apprill A, McNally S, Parsons R, Weber L. Minor revision to V4 region SSU rRNA 806R gene primer greatly increases detection of SAR11 bacterioplankton. Aquat Microb Ecol. 2015; 75:129–37.

31. Hildebrand F, Tadeo R, Voigt A, Bork P, Raes J. LotuS: an efficient and user-friendly OTU processing pipeline. Microbiome. 2014; 2:30.

32. Author. entropart: An R package to measure and partition diversity. Journal. 2015; doi: 10.18637/jss.v067.i08.

33. Oksanen J, Blanchet GF, Friendly M, Kindt R, Legendre P, McGilinn D, et al. Vegan: community ecology package. R package version 2.5-5. 2019. https://CRAN.R-project.org/package=vegan

34. R Core Team. R: A language and environment for statistical computing, reference index version 3.6.0. 2019. http://www.R-project.org

35. Albertsen M, Hugenholtz P, Skarshewski A, Nielsen KL, Tyson GW, Nielsen PH. Genome sequences of rare, uncultured bacteria obtained by differential coverage binning of multiple metagenomes. Nat Biotechnol. 2013; 31:533–8.

36. Rodriguez-R LM, Konstantinidis KT. Nonpareil: a redundancy-based approach to assess the level of coverage in metagenomic datasets. Bioinformatics. 2014; 30:629–35.

37. Rodriguez-R LM, Konstantinidis KT. Estimating coverage in metagenomic data sets and why it matters. ISME J. 2014; 8:2349–51.

38. Chevreux B, Pfisterer T, Drescher B, Driesel AJ, Müller WEG, Wetter T, et al. Using the miraEST Assembler for Reliable and Automated mRNA Transcript Assembly and SNP Detection in Sequenced ESTs. Genome Res. 2004; 14:1147–59.

39. Li D, Liu C-M, Luo R, Sadakane K, Lam T-W. MEGAHIT: an ultra-fast single-node solution for large and complex metagenomics assembly via succinct de Bruijn graph. Bioinformatics. 2015;31:1674–6.

40. Strous M, Kraft B, Bisdorf R, Tegetmeyer H. The binning of metagenomic contigs for microbial physiology of mixed cultures. Front Microbiol. 2012; 3:A410.

41. Rodriguez-R LM, Gunturu S, Harvey WT, Rosselló-Mora R, Tiedje JM, Cole JR, et al. The Microbial Genomes Atlas (MiGA) webserver: taxonomic and gene diversity analysis of Archaea and Bacteria at the whole genome level. Nucleic Acids Res. 2018; 46:W282–W8.

42. Seemann T. Prokka: rapid prokaryotic genome annotation. Bioinformatics. 2014; 30:2068–9.

43. Author. AromaDeg, a novel database for phylogenomics of aerobic bacterial degradation of aromatics. Journal. 2014; doi: 10.1093/database/bau118.

44. Leplae R, Lima-Mendez G, Toussaint A. ACLAME: A CLAssification of Mobile genetic Elements, update 2010. Nucleic Acids Res. 2010; 38:D57–D61.

45. Overbeek R, Olson R, Pusch GD, Olsen GJ, Davis JJ, Disz T, et al. The SEED and the Rapid Annotation of microbial genomes using Subsystems Technology (RAST). Nucleic Acids Res. 2014; 42:D206–D14.

46. Zhao Y, Tang H, Ye Y. RAPSearch2: a fast and memory-efficient protein similarity search tool for next-generation sequencing data. Bioinformatics. 2012; 28:125–6.

47. Guy L, Roat Kultima J, Andersson SGE. genoPlotR: comparative gene and genome visualization in R. Bioinformatics. 2010; 26:2334–5.

48. Dobin A, Davis CA, Schlesinger F, Drenkow J, Zaleski C, Jha S. STAR: ultrafast universal RNA-seq aligner. Bioinformatics. 2013; 29:15–29.

49. Anders S, Pyl PT, Huber W. HTSeq - a Python framework to work with high-throughput sequencing data. Bioinformatics. 2015; 31:166–9.

50. Robinson MD, Smyth GK. Moderated statistical tests for assessing differences in tag abundance. Bioinformatics. 2010; 23:2881–7.

51. Robinson MD, McCarthy DJ, Smyth GK. edgeR: a Bioconductor package for differential expression analysis of digital gene expression data. Bioinformatics. 2010; 26:139–40.

52. Lund SP, Nettleton D, McCarthy Davis J, Smyth Gordon K. Detecting differential expression in RNA-sequence data using quasi-likelihood with shrunken dispersion estimates. Statistical Applications in Genetics and Molecular Biology. 2012; 11:A8.

53. Clauset A, Newman MEJ, Moore C. Finding community structure in very large networks. Physical Review E. 2004; 70:066111.

54. Berry D, Widder S. Deciphering microbial interactions and detecting keystone species with co-occurrence networks. Front Microbiol. 2014; 5:219.

55. Csardi G, Nepusz T. The igraph software package for complex network research. InterJournal, Complex Systems. 2006; 1695:1695.

56. Seidel K, Kühnert J, Adrian L. The complexome of Dehalococcoides mccartyi reveals Its organohalide respiration-complex Is modular. Front Microbiol. 2018; 9:A1130.

57. Sirtori C, Agüera A, Carra I, Sanchéz Pérez JA. Identification and monitoring of thiabendazole transformation products in water during Fenton degradation by LC-QTOF-MS. Anal Bioanal Chem. 2014; 406:5323–37.

58. Campos-Mañas MC, Plaza-Bolaños P, Martínez-Piernas AB, Sánchez-Pérez JA, Agüera A. Determination of pesticide levels in wastewater from an agro-food industry: Target, suspect and transformation product analysis. Chemosphere. 2019; 232:152–63.

59. Martínez-Piernas AB, Plaza-Bolaños P, García-Gómez E, Fernández-Ibáñez P, Agüera A. Determination of organic microcontaminants in agricultural soils irrigated with reclaimed wastewater: Target and suspect approaches. Anal Chim Acta. 2018; 1030:115–24.

60. Horan K, Girke T, Backman TWH, Wang Y. fmcsR: mismatch tolerant maximum common substructure searching in R. Bioinformatics. 2013; 29:2792–4.

61. Chen AF, Cao D-S, Xiao N, Xu Q-S. Rcpi: R/Bioconductor package to generate various descriptors of proteins, compounds and their interactions. Bioinformatics. 2014; 31:279–81.

62. Waterhouse A, Bertoni M, Bienert S, Studer G, Tauriello G, Gumienny R, et al. SWISS-MODEL: homology modelling of protein structures and complexes. Nucleic Acids Res. 2018; 46:W296–W303.

63. Trott O, Olson AJ. AutoDock Vina: improving the speed and accuracy of docking with a new scoring function, efficient optimization, and multithreading. Journal of Computational Chemistry. 2010; 31:455–61.

64. Morris GM, Huey R, Lindstrom W, Sanner MF, Belew RK, Goodsell DS, et al. AutoDock4 and AutoDockTools4: Automated docking with selective receptor flexibility. Journal of Computational Chemistry. 2009; 30:2785–91.

65. Pettersen EF, Goddard TD, Huang CC, Couch GS, Greenblatt DM, Meng EC, et al. UCSF Chimera—A visualization system for exploratory research and analysis. Journal of Computational Chemistry. 2004; 25:1605–12.

66. Shintani M, Urata M, Inoue K, Eto K, Habe H, Omori T, et al. The *Sphingomonas* plasmid pCAR3 is involved in complete mineralization of carbazole. J Bacteriol. 2007; 189:2007–20.

67. Nojiri H. Structural and molecular genetic analyses of the bacterial carbazole degradation system. Biosci, Biotechnol, Biochem. 2012; 76:1–18.

68. Noinaj N, Guillier M, Barnard TJ, Buchanan SK. TonB-dependent transporters: regulation, structure, and function. Annu Rev Microbiol. 2010; 64:43–60.

69. Fang H, Li D, Kang J, Jiang P, Sun J, Zhang D. Metabolic engineering of *Escherichia coli* for de novo biosynthesis of vitamin B12. Nat Commun. 2018; 9:4917.

70. Author. Biosynthesis and use of cobalamin (B12). Journal. 2008; doi: doi:10.1128/ecosalplus.3.6.3.8.

71. Chimento DP, Mohanty AK, Kadner RJ, Wiener MC. Substrate-induced transmembrane signaling in the cobalamin transporter BtuB. Nat Struct Biol. 2003; 10:394–401.

72. Salam LB, Ilori MO, Amund OO. Properties, environmental fate and biodegradation of carbazole. 3 Biotech. 2017; 7:111.

73. Schmidt AW, Reddy KR, Knölker H-J. Occurrence, biogenesis, and synthesis of biologically active carbazole alkaloids. Chemical Reviews. 2012; 112:3193–328.

74. Gieg LM, Otter A, Fedorak PM. Carbazole degradation by *Pseudomonas* sp. LD2: metabolic characteristics and the identification of some metabolites. Environ Sci Technol. 1996; 30:575–85.

75. Schneider J, Grosser RJ, Jayasimhulu K, Xue W, Kinkle B, Warshawsky D. Biodegradation of carbazole by Ralstonia sp. RJGII.123 isolated from a hydrocarbon contaminated soil. Can J Microbiol. 2000; 46:269–77.

76. Habe H, Ashikawa Y, Saiki Y, Yoshida T, Nojiri H, Omori T. *Sphingomonas* sp. strain KA1, carrying a carbazole dioxygenase gene homologue, degrades chlorinated dibenzo-p-dioxins in soil. FEMS Microbiol Lett. 2002; 211:43–9.

77. Nam J-W, Nojiri H, Noguchi H, Uchimura H, Yoshida T, Habe H, et al. Purification and characterization of carbazole 1,9a-dioxygenase, a three-component dioxygenase system of *Pseudomonas* resinovorans strain CA10. Appl Environ Microbiol. 2002; 68:5882–90.

78. Gai Z, Wang X, Liu X, Tai C, Tang H, He X, et al. The genes coding for the conversion of carbazole to catechol are flanked by IS6100 elements in *Sphingomonas* sp. strain XLDN2-5. PLOS ONE. 2010; 5:e10018.

79. Kilbane Ii JJ, Daram A, Abbasian J, Kayser KJ. Isolation and characterization of Sphingomonas sp. GTIN11 capable of carbazole metabolism in petroleum. Biochem Biophys Res Commun. 2002; 297:242–8.

80. Sato SI, Nam JW, Kasuga K, Nojiri H, Yamane H, Omori T. Identification and characterization of genes encoding carbazole 1,9a-dioxygenase in *Pseudomonas* sp. strain CA10. J Bacteriol. 1997; 179:4850–8.

81. Inoue K, Habe H, Yamane H, Nojiri H. Characterization of novel carbazole catabolism genes from Gram-positive carbazole degrader *Nocardioides aromaticivorans* IC177. Appl Environ Microbiol. 2006; 72:3321–9.

82. Urata M, Uchimura H, Noguchi H, Sakaguchi T, Takemura T, Eto K, et al. Plasmid pCAR3 contains multiple gene sets involved in the conversion of carbazole to anthranilate. Appl Environ Microbiol. 2006; 72:3198–205.

83. Pinyakong O, Habe H, Yoshida T, Nojiri H, Omori T. Identification of three novel salicylate 1-hydroxylases involved in the phenanthrene degradation of Sphingobium sp. strain P2. Biochem Biophys Res Commun. 2003; 301:350–7.

84. Nojiri H, Nam J-W, Kosaka M, Morii K-I, Takemura T, Furihata K, et al. Diverse oxygenations catalyzed by carbazole 1,9a-dioxygenase from *Pseudomonas* sp. Strain CA10. J Bacteriol. 1999; 181:3105–13.

85. Aylward FO, McDonald BR, Adams SM, Valenzuela A, Schmidt RA, Goodwin LA, et al. Comparison of 26 Sphingomonad Genomes Reveals Diverse Environmental Adaptations and Biodegradative Capabilities. Appl Environ Microbiol. 2013; 79:3724–33.

86. Verma H, Kumar R, Oldach P, Sangwan N, Khurana JP, Gilbert JA, et al. Comparative genomic analysis of nine Sphingobium strains: insights into their evolution and hexachlorocyclohexane (HCH) degradation pathways. BMC Genomics. 2014; 15:1014.

87. Zhao Q, Yue S, Bilal M, Hu H, Wang W, Zhang X. Comparative genomic analysis of 26 Sphingomonas and Sphingobium strains: Dissemination of bioremediation capabilities, biodegradation potential and horizontal gene transfer. Sci Total Environ. 2017; 609:1238–47.

88. Stolz A. Degradative plasmids from sphingomonads. FEMS Microbiol Lett. 2014; 350:9–19.

89. Basta T, Keck A, Klein J, Stolz A. Detection and characterization of conjugative degradative plasmids in xenobiotic-degrading *Sphingomonas* strains. J Bacteriol. 2004; 186:3862–72.

90. Sangwan N, Xia FF, Gilbert JA. Recovering complete and draft population genomes from metagenome datasets. Microbiome. 2016; 4:A8.

91. Romine MF, Rodionov DA, Maezato Y, Osterman AL, Nelson WC. Underlying mechanisms for syntrophic metabolism of essential enzyme cofactors in microbial communities. ISME J. 2017; 11:1434.

92. Yan J, Im J, Yang Y, Löffler FE. Guided cobalamin biosynthesis supports *Dehalococcoides mccartyi* reductive dechlorination activity. Philosophical Transactions of the Royal Society B: Biological Sciences. 2013; 368:20120320.

93. Garcia SL, Buck M, McMahon KD, Grossart H-P, Eiler A, Warnecke F. Auxotrophy and intrapopulation complementary in the ‘interactome’ of a cultivated freshwater model community. Mol Ecol. 2015; 24:4449–59.

94. Payne KAP, Quezada CP, Fisher K, Dunstan MS, Collins FA, Sjuts H, et al. Reductive dehalogenase structure suggests a mechanism for B12-dependent dehalogenation. Nature. 2015; 517:513–6.

95. Wang S. Bacterial Two-Component Systems: Structures and Signaling Mechanisms. In: Huang C, editor. Protein Phosphorylation in Human Health: InTech; 2012.

96. Zschiedrich CP, Keidel V, Szurmant H. Molecular mechanisms of two-component signal transduction. J Mol Biol. 2016; 428:3752–75.

97. Schaefers MM, Liao TL, Boisvert NM, Roux D, Yoder-Himes D, Priebe GP. An oxygen-sensing two-component system in the *Burkholderia cepacia* complex regulates biofilm, intracellular invasion, and pathogenicity. PLoS Path. 2017; 13:e1006116.

98. Trastoy R, Manso T, Fernández-García L, Blasco L, Ambroa A, Pérez del Molino ML, et al. Mechanisms of bacterial tolerance and persistence in the gastrointestinal and respiratory environments. Clin Microbiol Rev. 2018; 31:e00023–18.

99. Green J, Stapleton MR, Smith LJ, Artymiuk PJ, Kahramanoglou C, Hunt DM, et al. Cyclic-AMP and bacterial cyclic-AMP receptor proteins revisited: adaptation for different ecological niches. Curr Opin Microbiol. 2014; 18:1–7.

100. Lin H, Hoffmann F, Rozkov A, Enfors S-O, Rinas U, Neubauer P. Change of extracellular cAMP concentration is a sensitive reporter for bacterial fitness in high-cell-density cultures of *Escherichia coli*. Biotechnol Bioeng. 2004; 87:602–13.

101. Slayden RA, Knudson DL, Belisle JT. Identification of cell cycle regulators in Mycobacterium tuberculosis by inhibition of septum formation and global transcriptional analysis. Microbiology. 2006; 152:1789–97.

102. Sarcina M, Mullineaux CW. Effects of tubulin assembly inhibitors on cell division in prokaryotes in vivo. FEMS Microbiol Lett. 2000; 191:25–9.

103. Kumar K, Awasthi D, Berger WT, Tonge PJ, Slayden RA, Ojima I. Discovery of anti-TB agents that target the cell-division protein FtsZ. Future Medicinal Chemistry. 2010; 2:1305–23.

104. Bers K, Leroy B, Breugelmans P, Albers P, Lavigne R, Sørensen SR, et al. A novel hydrolase identified by genomic-proteomic analysis of phenylurea herbicide mineralization by *Variovorax* sp. strain SRS16. Appl Environ Microbiol. 2011; 77:8754–64.

105. Vandera E, Samiotaki M, Parapouli M, Panayotou G, Koukkou AI. Comparative proteomic analysis of *Arthrobacter phenanthrenivorans* Sphe3 on phenanthrene, phthalate and glucose. Journal of Proteomics. 2015; 113:73–89.

106. Perruchon C, Vasileiadis S, Rousidou K, Papadopoulou E, Tanou G, Samiotaki M, et al. Metabolic pathway and cell adaptation mechanisms revealed through genomic, proteomic and transcription analysis of a Sphingomonas haloaromaticamans strain degrading ortho-phenylphenol. Scientific Reports. 2017; 7:6449.

107. Ferrer A, Rivera J, Zapata C, Norambuena J, Sandoval Á, Chávez R, et al. Cobalamin protection against oxidative stress in the acidophilic iron-oxidizing bacterium *Leptospirillum* group II CF-1. Front Microbiol. 2016; 7.

108. Yoon K-S, Tsukada N, Sakai Y, Ishii M, Igarashi Y, Nishihara H. Isolation and characterization of a new facultatively autotrophic hydrogen-oxidizing Betaproteobacterium, *Hydrogenophaga* sp. AH-24. FEMS Microbiol Lett. 2008; 278:94–100.

109. Morris JJ, Lenski RE, Zinser ER. The Black Queen Hypothesis: Evolution of Dependencies through Adaptive Gene Loss. mBio. 2012; 3:e00036–12.

