## Supplemental Material for "Nutritional inter-dependencies and a carbazole-dioxygenase are key elements of a bacterial consortium relying on a *Sphingomonas* for the degradation of the fungicide thiabendazole"

### Supporting Materials and Methods

**Media and culture conditions.** A minimal salts medium (MSMN) was used for the routine cultivation of the bacterial consortium. The MSMN consisted of (g $l^{-1}$): $KH_2PO_4$, (2.27), $Na_2HPO_4.12H_2O$ (5.97), $NH_4Cl$ (1.0), $MgSO_4.7H_2O$, (0.5) $CaCl_2.2H_2O$, (0.01), $MnSO_4.4H_2O$ (0.02) and $FeSO_4$, (0.025), with an overall pH of 6.8 All reagents for the MSMN were purchased from Sigma-Aldrich St. (Louis, USA). All solutions were autoclaved except for the iron-containing stock solution which was filter sterilized (0.22 μm, Q-max, Frisenette, Knebel, Denmark), and were aseptically added to the final medium. In all assays the inoculum was derived from fresh liquid cultures which had been amended with 25 μg $ml^{-1}$ TBZ (Sigma-Aldrich, St. Louis, Missouri, USA) as sole carbon source and was harvested when TBZ degradation was nearly complete, a point coinciding with maximum bacterial growth of the consortium as determined by preliminary q-PCR tests. TBZ was added to the medium to a final concentration of 25 μg $ml^{-1}$ (or 125 μM) from a filter-sterilized 125 mM DMSO stock solution. The final DMSO concentration in the medium was < 0.1 % with the same DMSO content being added in the TBZ free treatments. Filter sterilized succinate was added in equal carbon mass concentration (15 μg $ml^{-1}$ in each case) with the TBZ (37 μg $ml^{-1}$ final concentration; 314 μM). The TBZ used in the stable isotope experiments was uniformly $^{13}C$-labeled at its phenolic ring (purchased from Clearsynth®, Mumbai, India) (see Figure 1A).

**TBZ concentration measurement.** The concentration of TBZ in the growth medium during the degradation assays was monitored in aliquots of 0.5 ml which were regularly removed from the bacterial culture. TBZ residues were extracted by mixing the 0.5 ml of the aqueous medium with 1 ml methanol. The mixture was vortexed for 30 sec, centrifuged for 1 min at 17,000 xg and the clear supernatant was analyzed in a LabAlliance HPLC system equipped with a Marathon III pump (Rigas Labs, Thessaloniki, Greece) and a FASMA model 500 UV-Vis detector (Rigas Labs, Thessaloniki, Greece). The fungicide was eluted through a RP-C18 column (Athens, RP-C18, 120 Å, 4.6 mm x 150 mm, 5 μm) and it was detected at 254 nm using a mobile phase of 39% acetonitrile (ACN), 60.5% $H_2O$ and 0.5% $NH_3$. The concentration of TBZ was determined via external calibration curve prepared by injection of standard solutions of TBZ in methanol with concentrations ranging from 0.1 (limit of quantification) to 10 μg $ml^{-1}$.

**Nucleic acids extraction and 16S rRNA gene quantification.** DNA and RNA extractions were performed on bacterial cell pellets with the NucleoSpin® Tissue and RNA kits respectively (Macherey-Nagel & Co, Düren, Germany) according to the manufacturer's instructions. The nucleic acid extracts were quantified with the Quant-iT™ HS ds-DNA assay kit and the Quanti-iT™RNA HS kit with a Qubit™ fluorometer (Invitrogen, USA).

**Density gradient centrifugation and fractionation of labelled and non-labelled DNA.** DNA was extracted from bacterial cultures fed on $^{13}$C-labeled-TBZ and on unlabeled TBZ and quantified as described above. The DNA was separated into $^{13}$C-labeled and unlabeled ($^{12}$C) according to its molecular mass as described previously [1, 2]. Briefly, a calibration curve was generated with a density gradient generated by mixing a CsCl (≥99.999 % p.a., Carl Roth, Karlsruhe, Germany) solution (250 g of CsCl with 250 mL of sterile dd$H_2O$ water) with a gradient buffer (GBf - 50 ml of 1 M Tris-HCl, 3.75 g KCl and 1 ml of 0.5 M EDTA to 400 ml of water) at volume ratios starting from 100%/0% (CsCl/GBf) to 75%/25% (CsCl/GBf) by 5% volume increments. Each CsCl/GBf gradient was measured for its density by weight and its refraction index with an AR200 Handheld Digital refractometer (Reichert, Seefeld, Germany). The calculated calibration curve was used for setting the targeted initial CsCl density at 1.72 g ml$^{-1}$ for the DNA suspensions (using CsCl solution or GBf) prior ultracentrifugation, based on the refractive index values (1.4033 at 22 ˚C). The density adjusted DNA suspensions were transferred into 5.2 ml QuickSeal Polyallomer tubes (Beckman Coulter Pasadena, USA) of approximately equal weights, they were then heat sealed and centrifuged at 44,100 rpm (~167k x g), 20 ˚C, in vacuum for 36 hours in an Optima XPN-80 Ultracentrifuge (Beckman Coulter – maximum acceleration and minimum deceleration settings were employed) with the NVT 65.2 rotor (Beckman Coulter). The centrifuge tubes were carefully retrieved and secured at a stable position avoiding disturbing the generated CsCl gradient. The tubes were pierced near the top and right underneath the CsCl gradient surface with a syringe connected with a peristaltic pump through a tube supplied with constant sterile water flow. After, the tubes were pierced at the bottom and the pump was set to 1 ml min $^{-1}$ and activated, allowing the dripping of the tube content in a gradient-based order from the ultracentrifuge tube bottoms into 1.5 ml Eppendorf (Hamburg, Germany) tubes which were exchanged with empty ones (corresponding to the different density gradient fractions) every 25 seconds (~ 0.5 ml of CsCl gradient fractions per tube retrieved). The arrangement resulted in a constant transfer of the gradient in the 1.5 ml tubes and a stable liquid volume in the ultracentrifuge tubes. The refractive indexes of each collected CsCl-DNA suspension were measured and the subsample densities were calculated using the previously generated calibration curve. The DNA was purified from salts via glycogen/polyethylene glycol (PEG) precipitation as follows: One µl of glycogen and 2 volumes of PEG were added to the DNA suspensions. The tubes were mixed thoroughly, incubated at 4˚C for 15 hours and centrifuged at 13,000 g for 30 minutes at 15 ˚C. The supernatant was discarded and the pellet was washed with 500 µl of 70 % EtOH. After centrifugation for 10 minutes at 13,000 g and 15 ˚C, the supernatant was discarded, followed by drying in a desiccator at room temperature and in vacuum. The resulting pure DNA pellet was suspended in 30 µl of Tris-EDTA pH 8.0 and quantified as described above.

**16S rRNA gene diversity analysis.** DNA extracts were used for performing the 16S rRNA gene diversity analysis via PCR, during which the PCR products of all samples were indexed via a designed 5' extension of the forward primer, multiplexed in a single pool according to a previously setup protocol [3-5] and sequenced at a HiSeq2500 Illumina instrument. Briefly, the PCR reactions were performed into two steps as suggested by Berry, Mahfoudh [6] for avoiding primer index induced biases. In the first step 10 µl-PCR reactions were prepared using 2 µl of template DNA (normalized at 0.1 ng µl$^{-1}$), 0.4 µM of each primer (515f 5'-GTGYCAGCMGCCGCGGTAA-3', 806r 5'-GTGYCAGCMGCCGCGGTAA-3'),5 µl of the Q5® High Fidelity 2X master-mix (New England Biolabs®, Ipswich, MA, USA), 0.2 µl of molecular biology grade bovine serum albumin (BSA, New England Biolabs®, Ipswich, MA, USA), and volume was made upto 10 µl with PCR grade $H_2O$. The thermal cycling conditions were as follows: 3 minutes at 94 ˚C for enzyme activation, followed by 28 cycles of 30 seconds at 94 ˚C for denaturation, 30 seconds at 50 ˚C for annealing, and 30 seconds at 72 ˚C for extension, and with the cycles followed by 10 minutes at 72 ˚C of final extension. The second step PCR was performed in 20 µl reactions composed of 2 µl of the first step PCR product as template, 10 µl of the Q5® High Fidelity 2X master-mix, 0.4 µM of each primer (the reversed primer was the 806r while the forward was the 515f including the 5' linker index extension – complete list is provided in Table S2), and the rest was filled with PCR grade water. The thermal cycling conditions of the second step PCR were the same with those of the first step PCR with the sole difference that only 7 cycles (instead of 28) were performed. The second step PCR product concentrations were determined with the Qubit™ HS dsDNA assay kit (Invitrogen, Carlsbad, CA, USA) and equimolar amounts of each indexed product were multiplexed and purified with the Agencourt AMPureXP PCR purification kit (Beckman Coulter, Brea, CA, USA) prior to sequencing. The indexed primers were designed and selected according to their secondary formation with the barcrawl v100310 software [7].

Sequencing was performed at the Genome sequencing centre of the Brigham Young University (Provo, UT, USA) at a HiSeq2500 instrument at Rapid Mode generating 250 bp paired-end reads. The resulting sequences were quality controlled with Trimmomatic v0.36 [8] and the read pairs were demultiplexed according to their samples of origin (no index/primer mismatch allowed) with Flexbar v3.0[9] and assembled to their original amplicons, where possible, with the FLASH v1.2.8 software [10] for a minimum overlap of 40 bp. The sequences with more than one identical copy were processed with the Lotus v1.58 suit [11] using: UCHIME2 v4.2 [12] for the removal of chimeric sequences; USEARCH v10.0.240 for the 97 % identity operational taxonomic unit (OTU) calling; and the last common ancestor approach for sequence taxonomy annotation (acceptance of the 90% of the common lowest existing level of annotation of the best 20 hits) using BLAST v2.6.0+[13] search against the SILVA v132 database [14]. The resulting OTU and matrices were analysed for β-diversity shifts and differential abundance of OTUs/taxa between different conditions/time-points.

**Metagenome assembly, contig binning to genomes, annotation and synteny analysis.** Metagenome assembly was performed using the sequencing data of 5 shotgun libraries over three sequencing runs: one MiSeq Illumina (Illumina Inc., SanDiego, CA, USA) 300 bp paired-end reads run with the v3 kit reagents performed at the MrDNA facilities (Shallowater, TX, USA); one PacBio® RSII run (Pacific Biosciences, Menlo Park, CA, USA) with the P5-C3 (polymerase-chemistry versions) carried out at the McGill University and Génome Québec (Montréal, Québec, Canada) with the CCS reads being produced using the Quiver approach [15]; three Illumina HiSeq 2500, 250bp paired-end sequencing runs carried out with the Rapid SBS kit v2 chemistry at the GSC-BYU. The sequences were quality controlled, the Nonpareil algorithm [16, 17] was implemented (software version v3.301) in junction with the Nonpareil data analysis R package v3.3.1 [18] on the quality controlled sequences for assessing the achieved coverage given the sequencing effort. Hybrid assembly was performed with Mira v5. 1 [19] according to its quality scoring as part of the metAmos v1.5rc3 software suit [20]. The assembly was further improved with Megahit v1.1.3 [21] accepting k-mers with a minimum coverage cutoff of 5 and contigs of 500 bp minimum length, using concomitant sequence run outputs. Metawatt v3.5.3 [22] was used for contig binning performance according to GC content, coverage difference of taxa based on sampling condition differences (four DNA extracts were used for Illumina sequencing: three collected at time points corresponding to 50 % degradation, 100 % degradation and 24 hours post 100% degradation of TBZ, plus the DNA extracts obtained from the heavy DNA fraction of a sample collected at a time point corresponding to 10 % degradation of the $^{13}$C-labeled TBZ), codon usage, phylogenetic markers, tetra-nucleotide frequencies. The output metagenome assembled genomes (MAGs) were further assessed for classification and quality scoring with MiGA against its registered NCBI genome collection [23]. Annotation of the sequences was performed with Prokka v1.12 [24] at the metagenome annotation mode, functional annotations prioritized towards the aromatic hydrocarbon degradation AromaDeg [25] and the mobile genetic elements ALCME v0.4 [26] protein databases before searching for the default core set of proteins and profiles of Prokka. The gene calling process was further curated according to the RNAseq sequence mapping information generated as described below and the partial or complete genes identified via this method were searched with the basic local alignment search tool (BLAST) v2.8.1+ [13] against the NCBI nucleotide database prior introduction in the resulting annotation files. Further comparison of the proteins was carried out against the SEED database [27], using Rapsearch v2.22 [28] for complementing the annotation and exploiting the associated hierarchical annotation scheme. BLAST was also used for identifying molecular anchors during comparative genomics and the GenoPlotR v0.8.9 [29] R package was used for generating the associated plots.

**RNA sequencing and data analysis.** RNA sequencing was performed at an Illumina HiSeq2500 instrument with the Rapid SBS kit v2 chemistry for 250bp paired-end reads and the KAPA stranded RNA-Seq Library Preparation kit (KAPABiosystems, Wilmington, MA, USA) at the Genome Sequencing Center of the Brigham Young University (Provo, Utah, USA). Sequences were quality controlled as described for the metagenome sequencing and were mapped against the reference metagenome assembly sequence with STAR v020201 [30]. Transcript copy counts were predicted with HTSeq v 0.9.1[31]. Data quality control, counts per million (CPM) reads transformation, the trimmed mean of M-values (TMM) normalization approach [32] and treatment/time wise differential expression tests were performed with the edgeR v3.14.0 package [33]of the R v3.3.3 software [34] using the negative binomial models and the generalized linear model quasi likelihood F-test [35]. Genes with less than 4 CPM in at least 2 samples were rejected from further analysis. Redundancy analysis (RDA) on the counts per million and Hellinger transformed matrix values was preferred for testing the sample-wide main experimental effects (treatment – succinate vs TBZ – and time) over canonical correspondence analysis as previously suggested [36]. The Hellinger transformation (square root of the relative abundances) [37] was performed and multivariate analyses were performed with the Vegan v2.4-4 R package [38].

Correlation network analysis using the Pearson correlation r and Spearman rank sum correlation ρ coefficients were performed on the CPM matrices. The network analysis was performed in order to identify network density based sub-structures along with gene expression-driven keystone bins. The latter was performed by averaging the keystoneness index values of each expressed gene throughout each bin, following the methods described for microbial community matrices byBerry and Widder [39] for assessing the per expressed gene keystoneness indices. Five proposed keystoneness indicators used were the: (1) degree: being the number of connections of each network node (gene) according to the correlation test (Spearman's rank correlation coefficient with P-values adjusted to provide the false discovery rates for multiple hypothesis testing); (2) indirect degree: being the number of secondorder connections (genes connected to the directly connected genes of the examined gene); (3) betweenness centrality: being the number of shortest paths of any two nodes of the network which pass through the analysed node; (4) closeness centrality: being the average shortest path of any two nodes in the network other than the analysed; (5) transitivity: being the number of the observed triangles between the analysed node and any two connected nodes divided by all possible node numbers. Spearman correlation tests (Benjamini-Hochberg [40] P-value correction, α of 0.05 and only correlations with coefficient $\rho > 0.5$ were considered) were performed for the expressed gene-set for preparing the networks. The undirected networks, the density based sub-networks [41] and the keystoneness indices were calculated with the Igraph v1.0.1 package [42] of the R software. The keystoneness indices were further analysed for variance between bins using ANOVA and the Tukey's honestly significant differences *post hoc* test (α of 0.05).

**Shotgun protein analysis.** Samples corresponding to those used in the RNA sequencing analysis were used for shotgun proteomics. In brief, 2 ml of each sample was centrifuged at 10,000 g for 30 min, the supernatant was discarded, the pellets were washed with 1 ml of 50 mM ammonium bicarbonate (Ambic) buffer via centrifuging at 10,000 g for 30 minutes and resuspending in 30 μl Ambic buffer. The cells were disrupted by three freeze-thawing cycles of liquid nitrogen dipping followed by incubation at 40 ˚C for 60 sand sonication for 30 s at 37 kHz. 2 μl of GapDH (code G5262, Sigma, Taufkirchen, Germany) 20 μg $ml^{-1}$ (in sterile ultrapure water – Milli-Q, Merk-Millipore, Burlington, Massachusetts, USA) were spiked in the samples as internal standards. 2 μl of 1M dithiothreitol (DTT in 100 mM Ambic buffer) was applied (62.5 mM final concentration) and the samples were incubated at 30 ˚C for 1 h to reduce disulfide bonds. To prevent the reformation of disulfide bonds cysteine residues were derivatized with 15 μl of 400 mM 2-iodacetamide (IAA – 128 mM final concentration) and the samples were incubated at room temperature in the dark. Proteins were digested by using 6.3 μl of 0.1 μg $μl^{-1}$ trypsin stock solution in 1 mM HCl (0.63 μg of trypsin per sample – code T8658, Sigma, Taufkirchen, Germany), overnight at 37 ˚C. The digestion was stopped with the addition of 1 µl 100% formic acid per sample with glass pipettes and the digested peptides were obtained by retaining the supernatant after centrifugation at 16,000 g for 10 minutes.

The supernatants were dried in a Savant SPD1010 SpeedVac Concentrator (Thermo Fisher Scientific, Waltham, Massachusetts, USA) at room temperature to the volume of 10-20 µl and were desalted via a $ZipTip_{\mu-C18}$, (Merck-Millipore, Burlington, Massachusetts, USA), 2 µg capacity micro-column as follows. Five solutions were prepared containing 0%, 30%, 50%, 80%, 100% acetonitrile in $ddH_2O$ with the first 4 containing 0.1% formic acid. The micro-columns were washed and equilibrated by serially aspirating-dispensing 10 µl X 3 times each solution (last solution 5 times) following a descending order of acetonitrile concentration. Then, 10 µl of the sample was pipetted, washed by aspirating and discarding 10 µl of the 0 % acetonitrile solution, and collected the proteins into two collection tubes, one with the 30 % and one with 80 % acetonitrile solution by aspirating-dispensing 5 times in each tube. The pipetting procedure was performed three times for each sample and eventually the contents of the two collection tubes per sample were combined, dried in the Savant SpeedVac instrument and stored in -20 ˚C until analysis. For analysis, the dried and frozen samples were suspended in 20 µl 0.1 % formic acid in water by mixing for 10 min at room temperature before they were transferred to LC-MS sample vials and analysed with a nanoLC-coupled Thermo Orbitrap Fusion mass spectrometer (Thermo Fisher Scientific, Waltham, Massachusetts, USA) via a TriVersaNanoMate (Advion, Ltd., Harlow, UK). The chromatographic separation of peptides, parameters of electron spray ionization and mass spectrometry were as described previously[43]. Protein identification was done with Proteome Discoverer v2.2 (ThermoFisher Scientific, Waltham, MA, USA) with SequestHT as search engine [43] against the metagenome predicted proteins based on the metagenome annotated open reading frames (ORFs).

The abundances of the confidently predicted proteins were used as inputs for the data analysis in a similar manner with the RNA sequencing data (except for the network analysis), with the difference that no gene length-based correction approach was performed during data normalization.

**TBZ transformation product analysis and prediction.** Samples (4 ml) were collected from the growing cultures of the bacterial consortium fed on TBZ at regular intervals and stored at -80 ˚C until downstream analysis with LC-MS/MS. Prior to this the samples were thawed and they were subjected cell lysis by four sonication cycles at 80 kHz for 30 s. Then, the samples were filtered through 0.22-µm PTFE syringe filters. An aliquot of 450 µL was diluted with 25 µL of acetonitrile and 25 µL of a $^{13}C$-caffeine standard solution (Sigma-Aldrich, Steinheim, Germany), which was used as injection standard. The samples were then injected in an LC coupled to a quadrupole-time-of-flight mass analyser (LC-QTOF-MS). The ESI ionization source was set as follows: ion spray voltage, 4500 V; curtain gas, 25 (arbitrary units); GS1, 60 psi; GS2, 60 psi; and temperature, 575 °C. $N_2$was used as nebulizer, curtain and collision gas. A TOF MS survey scan was acquired followed by four IDA (Information Dependent Acquisition) TOF MS/MS scans within a *m/z* range from 100 to 2000 at a mass resolving power of 30000. The accumulation time applied was 250 ms and 100 ms for a TOF and an IDA scan, respectively. IDA criteria considered dynamic background subtraction. Collision energy of 30 eV with a ±15 eV spread was used in MS/MS fragmentation.

For the determination of TBZ transformation products, the conditions described in a previous work by Sirtori, Agüera [44], were applied. Acetonitrile (0.1 % formic acid and 5 % of water) and water (0.1 % formic acid) were used as eluents A and B, respectively. Water LC-MS and acetonitrile LC-MS grade were purchased from Honeywell (Seelze, Germany). The elution gradient was as follows: 10% A (3 min) to 100 % A in 22 min, which was hold for 3 min. The flow rate was 0.5 mL/min and the injection volume was 10 µL.

After the acquisition of the data by LC-QTOF-MS, data was processed using the strategy reported previously [45, 46]. The TBZ metabolite list was built using our approach to predict potential TBZ transformation products in the tested system. First, the SMILES [47] structural representations of TBZ were used for predicting compound transformants with SyGMa v1.1.0 according to the modelled phase 1 and 2 metabolism in the human body [48]. Following, the TBZ spectra were retrieved from the National Institute of Standards and Technology (NIST) of the US Department of Commerce website (http://webbook.nist.gov, last visited in 19 Dec. 2017) and were used for obtaining substructures with the MEtaboliteSubStructure Auto-Recommender (MESSAR) previously published by Mrzic, Meysman [49] freely available at http://messar.biodatamining.be (visited at 19 Dec 2017) and possible structural conformations with the MetfRag v2.4.2 [50] package as implemented in R v3.3.3[34]. Finally, MarvinSketch v17.28 (yr2017, ChemAxon http://www.chemaxon.com) of the Marvin suite was used for manually obtaining literature retrieved along with suspected metabolite SMILES representations. These resulted in 764 structural and 220 formula non redundant metabolites of the parent compound after screening according to InChI [51] and chemical formula respectively, with R. The Open-Babel v2.4.1 software suit [52] was used for structural format conversions and molecule associated property calculations.

In parallel with the above database construction, the rule-based prediction of the possible metabolites was carried out through the Eawag (Swiss Federal Institute of Aquatic Science and Technology) Biocatalysis/Biodegradation Database and Pathway Prediction System (EWAG-BBD/PPS; http://eawag-bbd.ethz.ch) whose rights were owned by the University of Minnesota (UM) until 2014 and is based on the UM-BBD/PPS search engine [53]. Another resource that was tested was the also web-based EnviPath pathway prediction package [54] but EWAG-BBD/PPS provided a more complete prediction.

**Carbazole/TBZ structural comparisons and *in silico* prediction of carbazole dioxygenase-TBZ interactions.** The up-regulated loci of carbazole dioxygenase in the *Sphingomonas* MAG 3X21F, with a putative role in the transformation of TBZ, were computationally analyzed for possible interaction prognosis with TBZ as a potential substrate. We initially assessed the similarity of the original substrate of these enzymes (i.e. carbazole) with TBZ, then the structural similarities of putative characterized homologues with our predicted enzymes, and finally the potential affinities of the encoded key enzyme with the target chemicals. Maximum common substructures between carbazole (original substrate of carbazole dioxygenase) and TBZ (relevant novel substrate of carbazole dioxygenase in our study) were calculated with the fmcsR v1.24.0 [55] R software package as implemented by Rcpi v1.18.1 [56] R software package. The protein three-dimensional structure models were calculated using SWISS-MODEL homology-based structure prediction approach [57]. Docking of TBZ was performed using Autodock Vina v1.1.2[58] and the Autodock Tools v4.2.6[59], while Chimera v1.11.2 [60] was used for further visualization and figure generation of the structures. The docking analysis preparation included the removal of non-polar hydrogens, the addition of the Kollman United Atom charges and the addition of the iron charges (found in the active sites) for the proteins, and the addition of the Gasteiger charges followed by the removal of the non-polar hydrogens for the ligands [61-63]. In the case of the flexible docking, residues around the active site were selected for allowing associated rotations and docking simulations were run. The Protein Data Bank [64] was used for protein structure searching, PubChem [65] was searched for ligand data, the Open-Babel v2.4.1 software suit [52] was used for structural format conversions and molecule associated property calculations, and the UGENE Unipro v1.32.0 [66] platform was used for the gene locus analysis.

### Supporting Figures

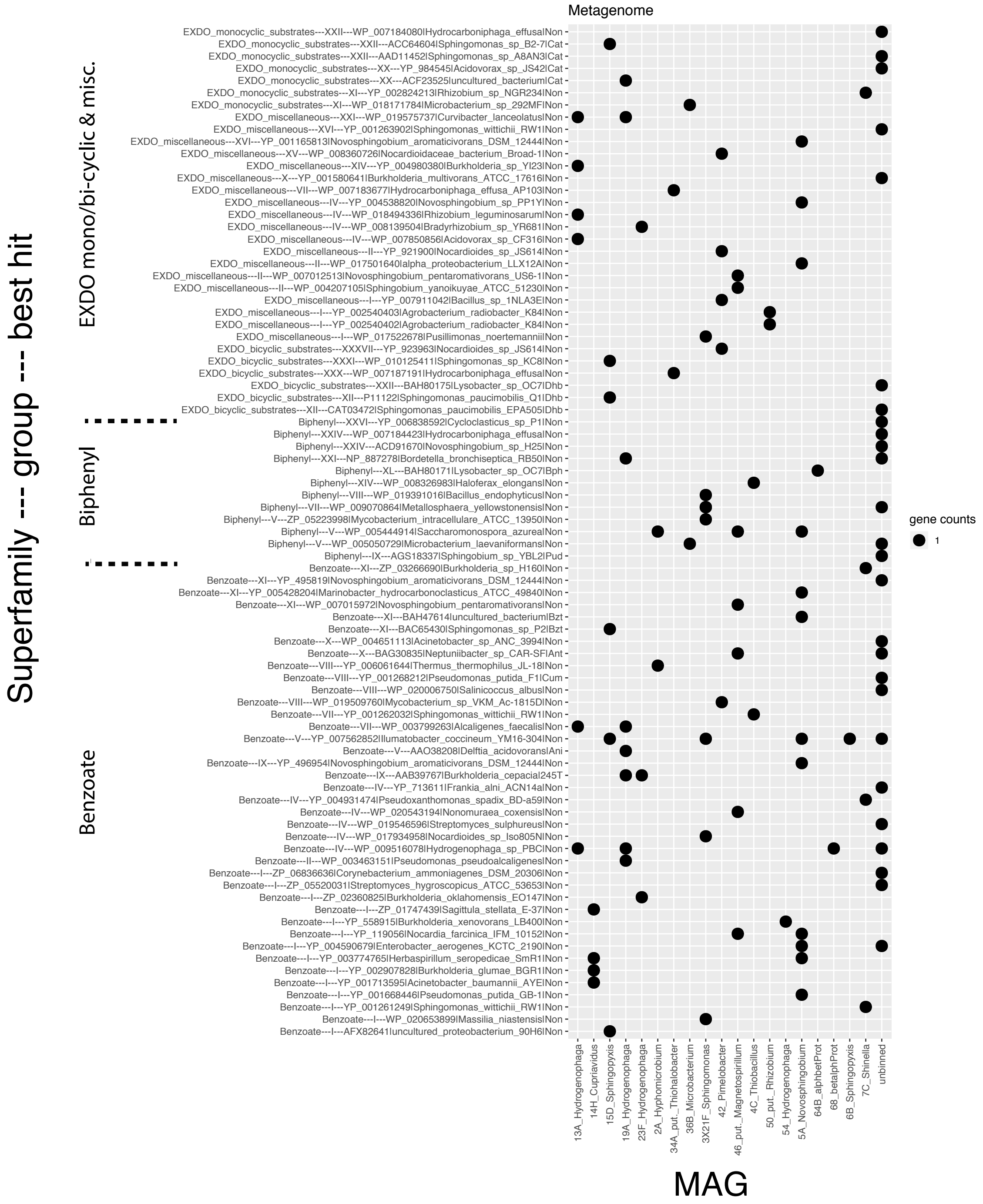

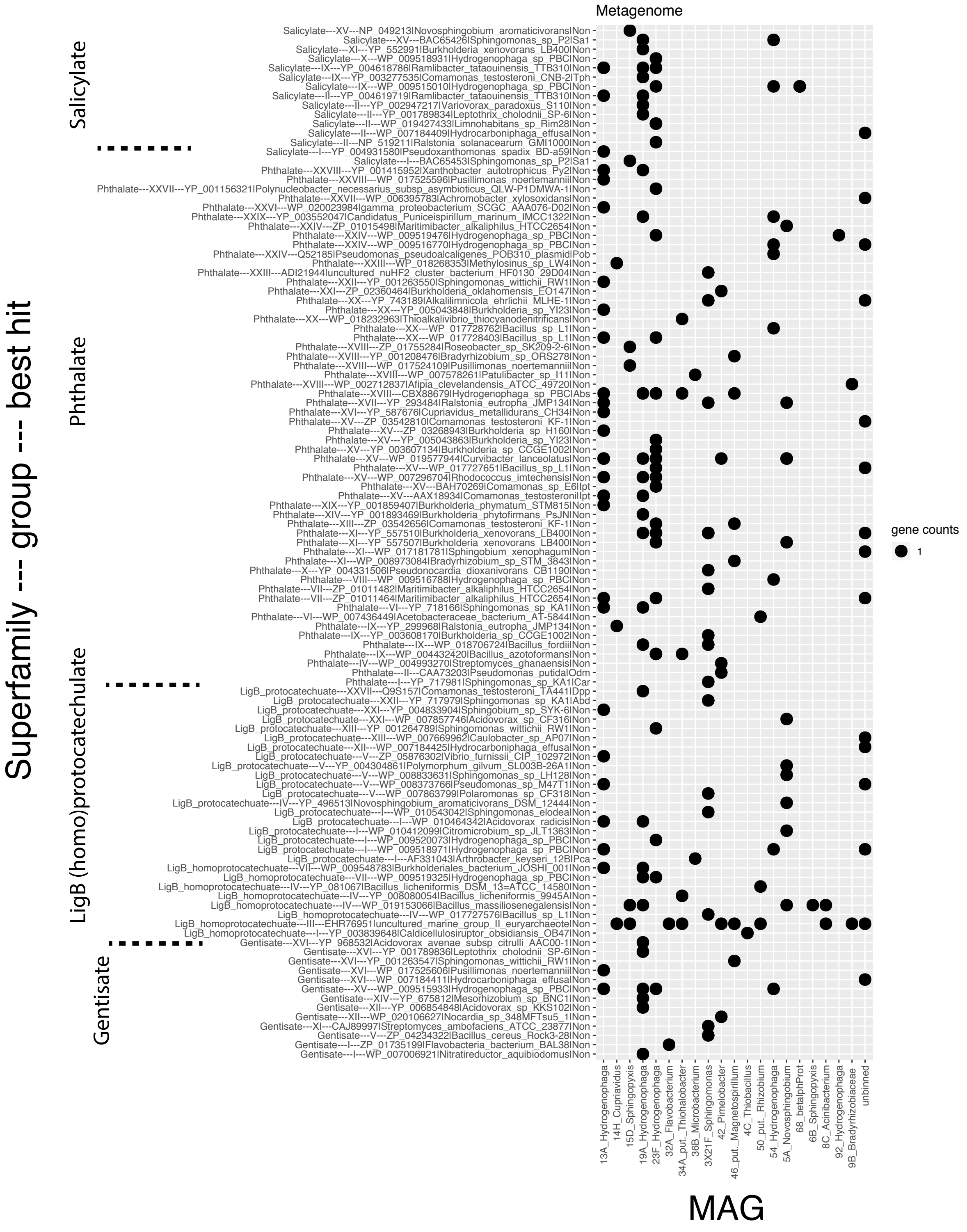

**Figure S1.** Bubble plot (presented in two pages) of the best metagenome translated open reading frame (ORF) BLAST hits (e-value cutoff of $1^{-10}$) on the AromaDeg database [25]. Assignment of the hits in the different metagenome assembled genomes (MAGs, provided as X-axis labels) is provided according to the key on the right. The enzyme superfamilies along with the phylogenetic clusters/groups and the best hits (including the hit accession numbers, owning organisms and experimentally verified substrate abbreviations after the pipe symbol "|") are provided as row labels.

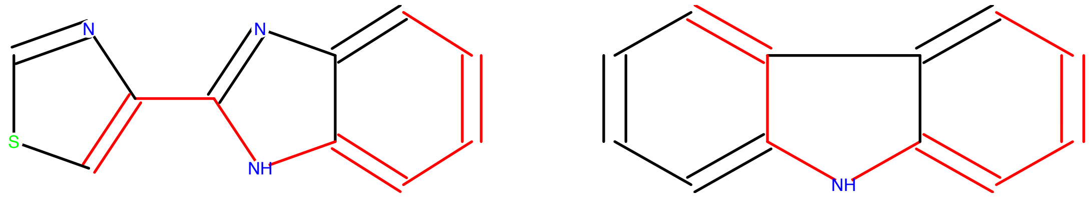


**Figure S2.** Common 3D substructures between thiabendazole (TBZ) (left) and carbazole (right) are highlighted with red. Concerning 3D structural similarity, we identified an overall molecule Tanimoto coefficient of 0.5 and a structural overlap index of 0.69.

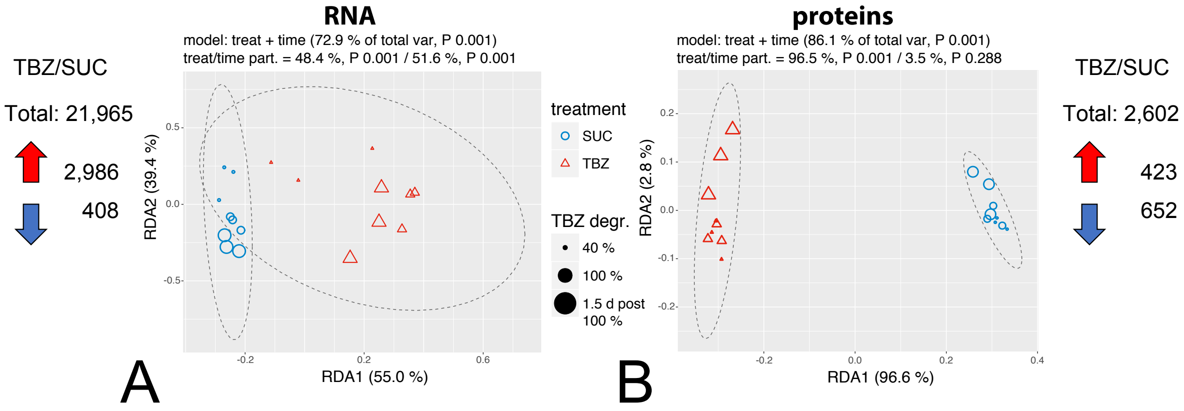


**Figure S3.** Redundancy analysis (RDA) of the gene expression profiles of the samples at RNA (A) and protein (B) level using the differentially expressed genes. Total expressed and differentially expressed genes (up-regulated or down-regulated at the TBZ compared with the succinate - SUC - treatment are depicted by the arrows) per molecule type analyzed are provided next to the RDA plots. Point sizes and colors are explained by the keys in between the two RDA plots, while tested model formulas, coinciding model variance and model significance along with their partition among tested variables are provided above each RDA plot.

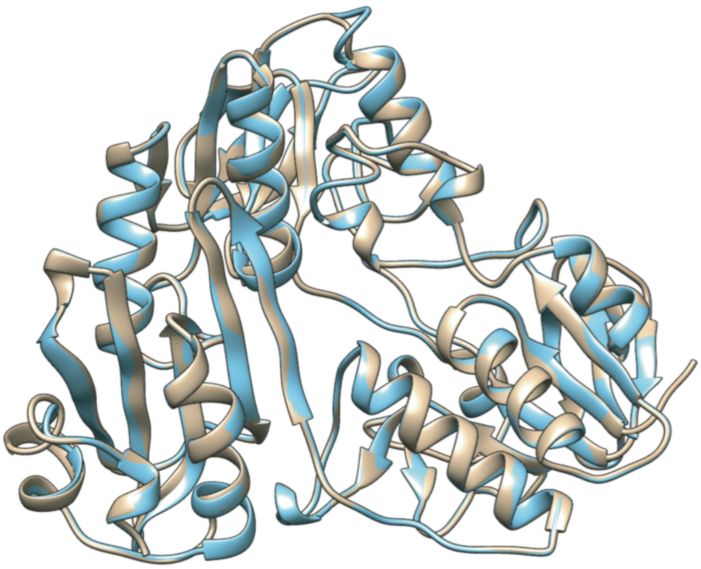

**Figure S4.** Structural comparison of the putative ferredoxin of the MAG 3X21F with locus tag number 00721 (found on QFCS01001167.1) against the *Novosphingobium* KA1 pCAR3 FdrI with acc. number BAF03314.1. Brown colors are used in predicted structures derived from our own consortium while the blue colors are used for ferredoxin residing on pCAR3. Due to the lack of crustal forms of any of the analyzed ferredoxins, the PDB accession 3LXD *Novosphingobium aromaticavorans* ferredoxin was used in both cases with SWISSMODEL as templates, being the closest PDB hits. The pCAR3 FdrI returned as best BLASTp hit the 00721 locus tag of MAG 3X21F (*Sphingomonas*) with statistics: Identities = 281/407 (69%), Positives = 324/407 (80%), Gaps = 0/407 (0%).

```
Identities = 285/369 (77%), Positives = 322/369 (87%), Gaps = 0/369 (0%)

Query  7    IAERRTKVWEPYIRAKLGFRNHWYPVRLASEIAEGTPVPVKLLGEKILLNRVGGKVYAIQ  66
            IAERRT+ W PYI AKLGFRNHWYPVRL++E+AE +PVPV+LLGEK+LLNRV G V+AI
Sbjct  10   IAERRTRAWAPYIDAKLGFRNHWYPVRLSAEVAEASPVPVQLLGEKVLLNRVDGVVHAIA  69

Query  67   DRCLHRGVTLSDRVECYSKNTISCWYHGWTYRWDDGRLVDILTNPGSVQIGRRALKTFPV  126
            DRCLHRGVTLSD+VECYSK TISCWYHGWTYRWD+G+LVDILTNP SVQIGR ALKT+PV
Sbjct  70   DRCLHRGVTLSDKVECYSKATISCWYHGWTYRWDNGKLVDILTNPTSVQIGRHALKTYPV  129

Query  127  EEAKGLIFVYVGDGEPTPLIEDVPPGFLDENRAIHGQHRLVASNWRLGAENGFDAGHVFI  186
             E KGL+F++VGD EP  L EDVPPGFLD + A+HGQHR+V +NWR+G ENGFDAGHVFI
Sbjct  130  REEKGLVFLFVGDQEPHDLAEDVPPGFLDADLAVHGQHRVVDANWRMGVENGFDAGHVFI  189

Query  187HKNSILVKGNDIILPLGFAPGDPDQLTRSEVAAGKPKGVYDLLGEHSVPVFEGMIEGKPA  246
HK+SIL+ GNDI LPLGFAPGDP+QLTRS    G PKGV+DLLGEHSVP+FE  IEG+PA
Sbjct  190HKSSILLDGNDIALPLGFAPGDPEQLTRSVTGEGAPKGVFDLLGEHSVPIFEATIEGQPA  249

Query  247  IHGNIGSKRVAISISIWLPGVLKVEPWPDPELTQFEWYVPVDETSHLYFQTLGKVVTSKE  306
            I G++GSK VAISIS+WLPGVLKV+P+PDP LTQFEWYVP+DE  HLY Q LG+ V S+E
Sbjct  250  IQGHMGSKMVAISISVWLPGVLKVDPFPDPTLTQFEWYVPIDEGHHLYLQMLGRRVGSEE  309

Query  307  AADSFEREFHEKWVGLALNGFNDDDIMARESMEPFYADDRGWSEEILFEPDRAIIEWRGL  366
A SFE EF EKWV LALNGFNDDDI+AR SMEPFYADDRGW EE+LFE DRAIIEWR L
Sbjct  310  EARSFEAEFREKWVELALNGFNDDDILARRSMEPFYADDRGWREEVLFESDRAIIEWRRL  369

Query  367  ASQHNRGIQ  375
            ASQ+NRGIQ
Sbjct  370  ASQYNRGIQ  378
```

**Figure S5**. Pairwise alignment (BLASTp [67]) of our own (Query) vs the *Novosphingobium* KA1 CarAa [68] (Subject). Active site residues including substrate interacting residues are shown in bold brown letters.

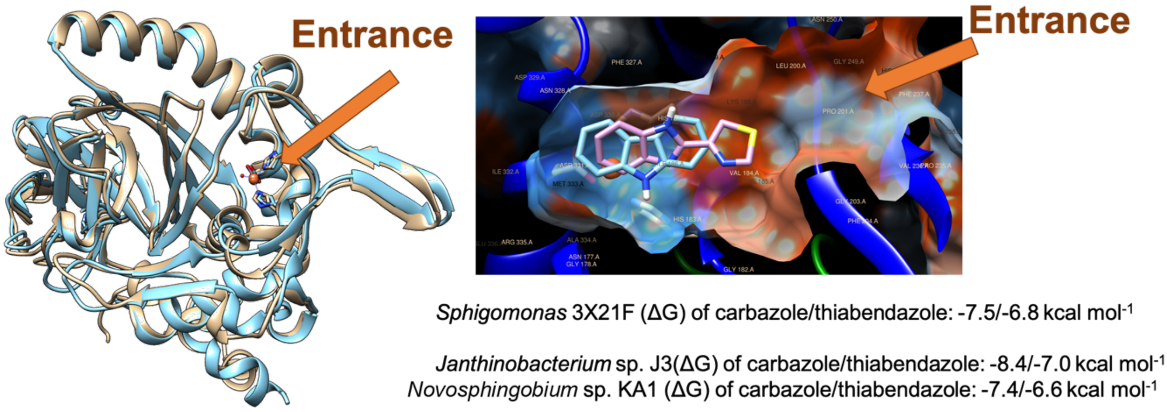


**Figure S6.**The 3D structure alignment of the CarAa homologue identified in the *car* operon of the *Sphingomonas* MAG 3X21F *vs* the *Novosphingobium* sp. KA1 1J1I protein databank (PDB) model (left) and the hydrophilic (blue)/hydrophobic (red) sites of the enzyme active center (right) together with the Gibbs free energy values for carbazole and thiabendazole docking analysis for *Sphingomonas* 3X21F CarAa, *Janthinobacterium* sp. J3 CarAa (PDB acc. 1WW9) and *Novosphingobium* sp. KA1 CarAa (PDB acc.3GKQ – also registered as *Novosphingobium* sp. KA1 CarAa).

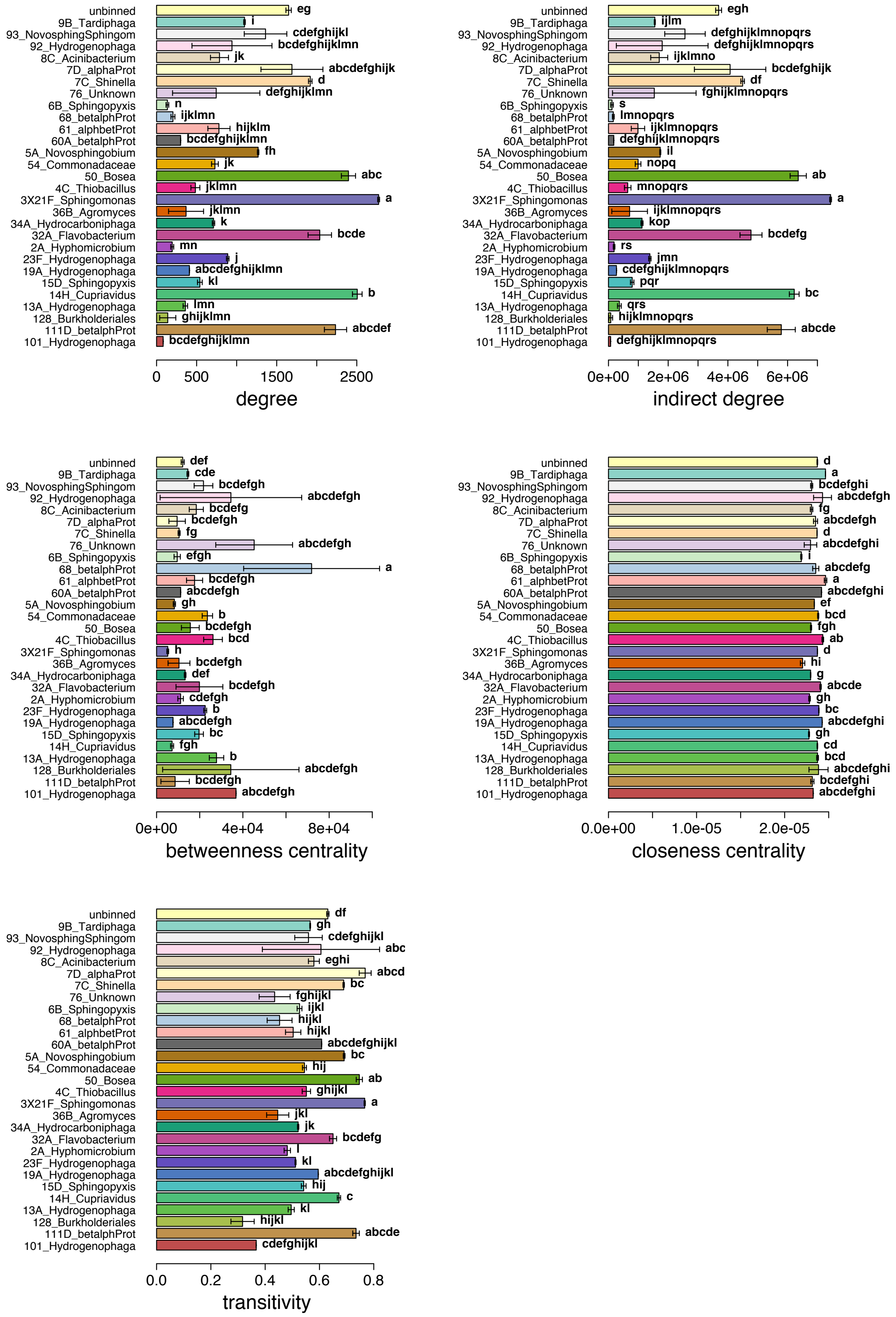


**Figure S7.** Keystoneness index barplots for the RNAseq participating MAGs according to the performed network analysis. Letters indicate significance groups according to the MAG related ANOVA and the Tukey post hoc analysis (αset to 0.05).

**Table S1.** Metagenome assembly stats (total number of features: 95542 genes; 94459 CDS; 6 ncRNA; 27 repeat_region (CRISPRs); 80 rRNA; 18 tmRNA; 978 tRNA). Metagenome Assembled Genomes (MAGs) smaller than 100 Kb were not included in the table and are collectively summarized at the bottom-left side of the table. The MAGs relative abundance columns provide information derived from samples collected at three time points corresponding to different TBZ degradation stages (50 % degradation, 100 % degradation and 24 hours after 100% degradation of TBZ).

| Metawatt bin IDs and stats | | | | | MiGA classification: NCBI hits | | | | MiGA: binning quality | | | MAG relative abundance | |
|---|---|---|---|---|---|---|---|---|---|---|---|---|---|
| **MAGID** | **contigs (#)** | **size (nt)** | **N50 (nt)** | **GC (%)** | **Phylum or Class** | **Strain** | **AAI or ANI (%)** | **Classification confidence level** | **Completeness (%)** | **Contamination (%)** | **Quality (%)** | **mean (%)** | **Standard deviation (%)** |
| 34A | 52 | 4,618,291 | 136,635 | 65.21 | γ-*Proteobacteria* | *Thiohalobacterthiocyanaticus* NZ AP018052 | 45.72% AAI | Class | 94.6 | 0.9 | 90.1 | 21.8 | 7.9 |
| 3X21F | 38 | 3,388,361 | 206,417 | 62.44 | α-*Proteobacteria* | *Sphingomonas* sp. DC 6 NZ CP021181 | 61.91% AAI | Genus | 95.5 | 0.9 | 91.0 | 17.9 | 7.1 |
| 9B | 25 | 4,220,254 | 315,044 | 62.13 | α-*Proteobacteria* | *Bradyrhizobiaceae* bacterium SG 6C NZ CM001195 | 92.33% ANI | Subspecies | 94.6 | 0.9 | 90.1 | 16.3 | 9.7 |
| 23F | 283 | 5,140,160 | 107,438 | 69.57 | β-*Proteobacteria* | *Hydrogenophaga* sp. PBC NZ CP017311 | 89.76% AAI | Subspecies | 89.2 | 2.7 | 75.7 | 8.9 | 4.2 |
| 19A | 189 | 5,487,813 | 61,470 | 65.83 | β-*Proteobacteria* | *Hydrogenophaga* sp. RAC07 | 78.96% AAI | Subspecies | 90.1 | 1.8 | 81.1 | 4.5 | 6.3 |
| 13A | 118 | 5,825,532 | 101,074 | 67.84 | β-*Proteobacteria* | *Hydrogenophaga* sp. RAC07 | 81.07% AAI | Subspecies | 91.9 | 6.3 | 60.4 | 3.2 | 1.8 |
| 4C | 50 | 3,365,496 | 183,522 | 62.47 | β-*Proteobacteria* | *Thiobacillusdenitrificans* ATCC 25259 NC 007404 | 78.5% AAI | Subspecies | 95.5 | 0.9 | 91.0 | 2.2 | 1.1 |
| 2A | 14 | 4,406,944 | 1,007,142 | 59.98 | α-*Proteobacteria* | *Hyphomicrobium* sp. MC1 NC 015717 | 76.37% AAI | Subspecies | 94.6 | 0.9 | 90.1 | 2.0 | 1.7 |
| 14H | 119 | 5,991,108 | 316,027 | 65.82 | α-*Proteobacteria* | *Mesorhizobiumamorphae* CCNWGS0123 NZ CP015318 | 64.33% AAI | Genus | 94.6 | 1.8 | 85.6 | 1.5 | 1.9 |
| 7C | 61 | 5,841,895 | 231,421 | 65.08 | α-*Proteobacteria* | *Shinella* sp. HZN7 NZ CP015736 | 91.07% ANI | Subspecies | 93.7 | 0.9 | 89.2 | 1.5 | 0.0 |
| 8C | 38 | 3,743,820 | 168,784 | 39.74 | *Bacteroidetes* | *Filimonas lacunae* NZ AP017422 | 60.87% AAI | Genus | 93.7 | 0.9 | 89.2 | 1.1 | 0.1 |
| 32A | 71 | 3,635,846 | 222,391 | 37.85 | *Bacteroidetes* | *Flavobacterium johnsoniae* NZ CP016907 | 64.1% AA | Genus | 95.5 | 0.9 | 91.0 | 0.9 | 1.2 |
| 5A | 21 | 4,388,512 | 636,638 | 66.38 | α-*Proteobacteria* | *Novosphingobiumaromaticivorans* DSM 12444 NC 007794T | 67.13% AAI | Genus | 95.5 | 0.0 | 95.5 | 0.6 | 0.5 |
| 54 | 784 | 2,974,980 | 101,474 | 68.86 | β-*Proteobacteria* | *Hydrogenophaga* sp. PBC NZ CP017311 | 98.65% ANI | Subspecies | 36.0 | 0.9 | 31.5 | 0.5 | 0.3 |
| 36B | 67 | 3,549,152 | 433,757 | 68.72 | *Actinobacteria* | *Microbacterium hominis* NZ CP025299 | 73.25% AAI | Subspecies | 89.2 | 0.9 | 84.7 | 0.5 | 0.4 |
| 42 | 792 | 4,947,347 | 36,913 | 72.63 | *Actinobacteria* | *Pimelobacter simplex* NZ CP009896 | 77.81% AAI | Subspecies | 86.5 | 3.6 | 68.5 | 0.4 | 0.6 |
| 15D | 88 | 4,025,560 | 283,945 | 66.11 | α-*Proteobacteria* | *Sphingopyxis* sp. WS5A3p CM009578 | 87.18% AAI | Subspecies | 92.8 | 2.7 | 79.3 | 0.5 | 0.1 |
| 50 | 1361 | 4,402,066 | 30,470 | 66.68 | α-*Proteobacteria* | *Rhizobium etli*8C 3 NZ CP017241 | 48.65% AAI | Order | 55.9 | 1.8 | 46.9 | 0.3 | 0.4 |
| 6B | 30 | 3,573,986 | 618,385 | 64.93 | α-*Proteobacteria* | *Sphingopyxis terrae* NBRC 15098 NZ CP013342 | 96.46% ANI | Subspecies | 93.7 | 1.8 | 84.7 | 0.3 | 0.0 |
| 46 | 366 | 6,274,208 | 86,277 | 66.48 | α-*Proteobacteria* | *Magnetospirillum* sp. ME 1 NZ CP015848 | 49.05% AAI | Order | 85.6 | 0.9 | 81.1 | 0.1 | 0.0 |

| | | | |
|---|---|---|---|
| <100Kb MAGs | 571 | 1,464,056 | 4,814 |
| Unbined contigs | 1650 | 7,278,568 | 24,154 |

| | | |
|---|---|---|
| **Total** | **6788** | **98,543,955** |

**Table S2.** The sequences of the barcoded primers used for the amplicon sequencing analysis of the 16S rRNA gene. The initial 9 bases comprise the index, followed by a GT linker and the 515F/806R primers [69, 70]. Treatment notation: "12C" for unlabeled thiabendazole in MSMN cultures; "13C" for $^{13}$C-labeled thiabendazole in MSMN cultures; "heavy" for the heavy CsCl gradient fraction retrieved DNA; and "light" for the light gradient fraction retrieved DNA.

| Sample ID | Barcoded Primer | Treatment | Time(hours post inoculation) |
|---|---|---|---|
| 01_12C2h36f9 | TTCTTCTTCGTGTGYCAGCMGCCGCGGTAA | 12C | 36 |
| 02_12C3h36f9 | TTCTCAATGGTGTGYCAGCMGCCGCGGTAA | 12C | 36 |
| 03_13C2h36f5 | TTCAGTTCAGTGTGYCAGCMGCCGCGGTAA | 13C_heavy | 36 |
| 04_13C2h36f9 | TTCGAATCAGTGTGYCAGCMGCCGCGGTAA | 13C_light | 36 |
| 05_13C3h36f5 | TTGTCAGGTGTGTGYCAGCMGCCGCGGTAA | 13C_heavy | 36 |
| 06_13C3h36f9 | TTGAAGTTCGTGTGYCAGCMGCCGCGGTAA | 13C_light | 36 |
| 07_12C1h72f9 | TTGCAACAAGTGTGYCAGCMGCCGCGGTAA | 12C | 72 |
| 08_12C2h72f9 | TTGGACGACGTGTGYCAGCMGCCGCGGTAA | 12C | 72 |
| 09_12C3h72f9 | TTCTTCAAGGTGTGYCAGCMGCCGCGGTAA | 12C | 72 |
| 10_13C1h72f5 | TTCTCAGAAGTGTGYCAGCMGCCGCGGTAA | 13C_heavy | 72 |
| 11_13C1h72f9 | TTCAGTAAGGTGTGYCAGCMGCCGCGGTAA | 13C_light | 72 |
| 12_13C2h72f5 | TTCGACAATGTGTGYCAGCMGCCGCGGTAA | 13C_heavy | 72 |
| 13_13C2h72f9 | TTGTCGATAGTGTGYCAGCMGCCGCGGTAA | 13C_light | 72 |
| 14_13C3h72f5 | TTGAAGGAAGTGTGYCAGCMGCCGCGGTAA | 13C_heavy | 72 |
| 15_13C3h72f9 | TTGCAGTATGTGTGYCAGCMGCCGCGGTAA | 13C_light | 72 |
| 16_12C1h117f9 | TATATCAGGGTGTGYCAGCMGCCGCGGTAA | 12C | 117 |
| 17_12C2h117f9 | TTCTTGTCAGTGTGYCAGCMGCCGCGGTAA | 12C | 117 |
| 18_12C3h117f9 | TTCATATGGGTGTGYCAGCMGCCGCGGTAA | 12C | 117 |
| 19_13C1h117f5 | TTCAGACTTGTGTGYCAGCMGCCGCGGTAA | 13C_heavy | 117 |
| 20_13C1h117f9 | TTCGAGCACGTGTGYCAGCMGCCGCGGTAA | 13C_light | 117 |
| 21_13C2h117f5 | TTGTGTATCGTGTGYCAGCMGCCGCGGTAA | 13C_heavy | 117 |
| 22_13C2h117f9 | TTGACTATGGTGTGYCAGCMGCCGCGGTAA | 13C_light | 117 |
| 23_13C3h117f5 | TTGCCTAGTGTGTGYCAGCMGCCGCGGTAA | 13C_heavy | 117 |
| 24_13C3h117f9 | TATATCGTCGTGTGYCAGCMGCCGCGGTAA | 13C_light | 117 |
| 25_12C1h141f9 | TTCTTGAGTGTGTGYCAGCMGCCGCGGTAA | 12C | 141 |
| 26_12C2h141f9 | TTCATAGTCGTGTGYCAGCMGCCGCGGTAA | 12C | 141 |
| 27_12C3h141f9 | TTCAGAGGAGTGTGYCAGCMGCCGCGGTAA | 12C | 141 |
| 28_13C1h141f5 | TTGTTCAGAGTGTGYCAGCMGCCGCGGTAA | 13C_heavy | 141 |
| 29_13C1h141f9 | TTGTGTGAAGTGTGYCAGCMGCCGCGGTAA | 13C_light | 141 |
| 30_13C2h141f5 | TTGACGTGAGTGTGYCAGCMGCCGCGGTAA | 13C_heavy | 141 |
| 31_13C2h141f9 | TTGCCTCACGTGTGYCAGCMGCCGCGGTAA | 13C_light | 141 |
| 32_13C3h141f5 | TATATGCACGTGTGYCAGCMGCCGCGGTAA | 13C_heavy | 141 |
| 33_13C3h141f9 | TTCTTGGACGTGTGYCAGCMGCCGCGGTAA | 13C_light | 141 |

**References**


1. Neufeld JD, Vohra J, Dumont MG, Lueders T, Manefield M, Friedrich MW, et al. DNA stable-isotope probing. Nat Protocols. 2007; 2:860-6.
2. Perruchon C, Chatzinotas A, Omirou M, Vasileiadis S, Menkissoglou-Spiroudi U, Karpouzas DG. Isolation of a bacterial consortium able to degrade the fungicide thiabendazole: the key role of a *Sphingomonas* phylotype. Appl Microbiol Biotechnol. 2017; 101:3881-93.
3. Garau G, Silvetti M, Vasileiadis S, Donner E, Diquattro S, Deiana S, et al. Use of municipal solid wastes for chemical and microbiological recovery of soils contaminated with metal(loid)s. Soil Biol Biochem. 2017; 111:25-35.
4. Soldi S, Vasileiadis S, Uggeri F, Campanale M, Morelli L, Fogli M, et al. Modulation of the gut microbiota composition by rifaximin in non-constipated irritable bowel syndrome patients: a molecular approach. Clinical and Experimental Gastroenterology. 2015; 8:309-25.
5. Vasileiadis S, Puglisi E, Trevisan M, Scheckel KG, Langdon KA, McLaughlin MJ, et al. Changes in soil bacterial communities and diversity in response to long-term silver exposure. FEMS Microbiol Ecol. 2015; 91:fiv114.
6. Berry D, Mahfoudh KB, Wagner M, Loy A. Barcoded primers used in multiplex amplicon pyrosequencing bias amplification. Appl Environ Microbiol. 2011; 77:7846-9.
7. Frank DN. BARCRAWL and BARTAB: software tools for the design and implementation of barcoded primers for highly multiplexed DNA sequencing. BMC Bioinformatics. 2009; 10:362.
8. Bolger AM, Lohse M, Usadel B. Trimmomatic: A flexible trimmer for Illumina Sequence Data. Bioinformatics. 2014; 30:2114-20.
9. Dodt M, Roehr JT, Ahmed R, Dieterich C. FLEXBAR—Flexible Barcode and Adapter Processing for Next-Generation Sequencing Platforms. Biology. 2012; 1:895-905.
10. Magoč T, Salzberg SL. FLASH: fast length adjustment of short reads to improve genome assemblies. Bioinformatics. 2011.
11. Hildebrand F, Tadeo R, Voigt A, Bork P, Raes J. LotuS: an efficient and user-friendly OTU processing pipeline. Microbiome. 2014; 2:30.
12. Edgar R. UCHIME2: improved chimera prediction for amplicon sequencing. bioRxiv. 2016.
13. Altschul S, Gish W, Miller W, Myers E, Lipman D. Basic local alignment search tool. J Mol Biol. 1990; 215:403 - 10.
14. Yilmaz P, Parfrey LW, Yarza P, Gerken J, Pruesse E, Quast C, et al. The SILVA and "All-species Living Tree Project (LTP)" taxonomic frameworks. Nucleic Acids Res. 2014; 42:D643-D8.
15. Chin C-S, Alexander DH, Marks P, Klammer AA, Drake J, Heiner C, et al. Nonhybrid, finished microbial genome assemblies from long-read SMRT sequencing data. Nat Methods. 2013; 10:563.
16. Rodriguez-R LM, Konstantinidis KT. Nonpareil: a redundancy-based approach to assess the level of coverage in metagenomic datasets. Bioinformatics. 2014; 30:629-35.
17. Rodriguez-R LM, Konstantinidis KT. Estimating coverage in metagenomic data sets and why it matters. ISME J. 2014; 8:2349-51.
18. Rodriguez-R LM. Nonpareil: metagenome coverage estimation and projections for 'Nonpareil'. R package version 3.3.1. 2018. https://CRAN.R-project.org/package=Nonpareil
19. Chevreux B, Pfisterer T, Drescher B, Driesel AJ, Müller WEG, Wetter T, et al. Using the miraEST Assembler for Reliable and Automated mRNA Transcript Assembly and SNP Detection in Sequenced ESTs. Genome Res. 2004; 14:1147-59.
20. Treangen T, Koren S, Sommer D, Liu B, Astrovskaya I, Ondov B, et al. MetAMOS: a modular and open source metagenomic assembly and analysis pipeline. Genome Biology. 2013; 14:R2.
21. Li D, Liu C-M, Luo R, Sadakane K, Lam T-W. MEGAHIT: an ultra-fast single-node solution for large and complex metagenomics assembly via succinct de Bruijn graph. Bioinformatics. 2015; 31:1674-6.
22. Strous M, Kraft B, Bisdorf R, Tegetmeyer H. The binning of metagenomic contigs for microbial physiology of mixed cultures. Front Microbiol. 2012; 3:A410.

23. Rodriguez-R LM, Gunturu S, Harvey WT, Rosselló-Mora R, Tiedje JM, Cole JR, et al. The Microbial Genomes Atlas (MiGA) webserver: taxonomic and gene diversity analysis of Archaea and Bacteria at the whole genome level. Nucleic Acids Res. 2018; 46:W282-W8.
24. Seemann T. Prokka: rapid prokaryotic genome annotation. Bioinformatics. 2014; 30:2068-9.
25. Author. AromaDeg, a novel database for phylogenomics of aerobic bacterial degradation of aromatics. Journal. 2014; doi: 10.1093/database/bau118.
26. Leplae R, Lima-Mendez G, Toussaint A. ACLAME: A CLAssification of Mobile genetic Elements, update 2010. Nucleic Acids Res. 2010; 38:D57-D61.
27. Overbeek R, Olson R, Pusch GD, Olsen GJ, Davis JJ, Disz T, et al. The SEED and the Rapid Annotation of microbial genomes using Subsystems Technology (RAST). Nucleic Acids Res. 2014; 42:D206-D14.
28. Zhao Y, Tang H, Ye Y. RAPSearch2: a fast and memory-efficient protein similarity search tool for next-generation sequencing data. Bioinformatics. 2012; 28:125-6.
29. Guy L, Roat Kultima J, Andersson SGE. genoPlotR: comparative gene and genome visualization in R. Bioinformatics. 2010; 26:2334-5.
30. Dobin A, Davis CA, Schlesinger F, Drenkow J, Zaleski C, Jha S. STAR: ultrafast universal RNA-seq aligner. Bioinformatics. 2013; 29:15-29.
31. Anders S, Pyl PT, Huber W. HTSeq - a Python framework to work with high-throughput sequencing data. Bioinformatics. 2015; 31:166–9.
32. Robinson MD, Smyth GK. Moderated statistical tests for assessing differences in tag abundance. Bioinformatics. 2010; 23:2881–7.
33. Robinson MD, McCarthy DJ, Smyth GK. edgeR: a Bioconductor package for differential expression analysis of digital gene expression data. Bioinformatics. 2010; 26:139-40.
34. R Core Team. R: A language and environment for statistical computing, reference index version 3.6.0. 2019. http://www.R-project.org
35. Lund SP, Nettleton D, McCarthy Davis J, Smyth Gordon K. Detecting differential expression in RNA-sequence data using quasi-likelihood with shrunken dispersion estimates. Statistical Applications in Genetics and Molecular Biology. 2012; 11:A8.
36. (eds). Multivariate analysis of ecological data using CANOCO. (Cambridge Press,New York, 2003)
37. Legendre P, Gallagher E. Ecologically meaningful transformations for ordination of species data. Oecologia. 2001; 129:271-80.
38. Oksanen J, Blanchet GF, Friendly M, Kindt R, Legendre P, McGilinn D, et al. Vegan: community ecology package. R package version 2.5-5. 2019. https://CRAN.R-project.org/package=vegan
39. Berry D, Widder S. Deciphering microbial interactions and detecting keystone species with co-occurrence networks. Front Microbiol. 2014; 5:219.
40. Benjamini Y, Hochberg Y. Controlling the False Discovery Rate: A Practical and Powerful Approach to Multiple Testing. Journal of the Royal Statistical Society Series B (Methodological). 1995; 57:289-300.
41. Clauset A, Newman MEJ, Moore C. Finding community structure in very large networks. Physical Review E. 2004; 70:066111.
42. Csardi G, Nepusz T. The igraph software package for complex network research. InterJournal, Complex Systems. 2006; 1695:1695.
43. Seidel K, Kühnert J, Adrian L. The complexome of Dehalococcoides mccartyi reveals Its organohalide respiration-complex Is modular. Front Microbiol. 2018; 9:A1130.
44. Sirtori C, Agüera A, Carra I, Sanchéz Pérez JA. Identification and monitoring of thiabendazole transformation products in water during Fenton degradation by LC-QTOF-MS. Anal Bioanal Chem. 2014; 406:5323-37.
45. Campos-Mañas MC, Plaza-Bolaños P, Martínez-Piernas AB, Sánchez-Pérez JA, Agüera A. Determination of pesticide levels in wastewater from an agro-food industry: Target, suspect and transformation product analysis. Chemosphere. 2019; 232:152-63.
46. Martínez-Piernas AB, Plaza-Bolaños P, García-Gómez E, Fernández-Ibáñez P, Agüera A. Determination of organic microcontaminants in agricultural soils irrigated with reclaimed wastewater: Target and suspect approaches. Anal Chim Acta. 2018; 1030:115-24.

47. Weininger D. SMILES, a chemical language and information system. 1. Introduction to methodology and encoding rules. J Chem Inf Comput Sci. 1988; 28.
48. Ridder L, Wagener M. SyGMa: Combining Expert Knowledge and Empirical Scoring in the Prediction of Metabolites. ChemMedChem. 2008; 3:821-32.
49. Mrzic A, Meysman P, Bittremieux W, Laukens K. Automated Recommendation Of Metabolite Substructures From Mass Spectra Using Frequent Pattern Mining. bioRxiv. 2017.
50. Ruttkies C, Schymanski EL, Wolf S, Hollender J, Neumann S. MetFrag relaunched: incorporating strategies beyond in silico fragmentation. Journal of Cheminformatics. 2016; 8:3.
51. Heller SR, McNaught A, Pletnev I, Stein S, Tchekhovskoi D. InChI, the IUPAC International Chemical Identifier. Journal of Cheminformatics. 2015; 7:23.
52. O'Boyle NM, Banck M, James CA, Morley C, Vandermeersch T, Hutchison GR. Open Babel: An open chemical toolbox. Journal of Cheminformatics. 2011; 3:33.
53. Gao J, Ellis LBM, Wackett LP. The University of Minnesota Biocatalysis/Biodegradation Database: improving public access. Nucleic Acids Res. 2010; 38:D488-D91.
54. Wicker J, Lorsbach T, Gütlein M, Schmid E, Latino D, Kramer S, et al. enviPath – The environmental contaminant biotransformation pathway resource. Nucleic Acids Res. 2016; 44:D502-D8.
55. Horan K, Girke T, Backman TWH, Wang Y. fmcsR: mismatch tolerant maximum common substructure searching in R. Bioinformatics. 2013; 29:2792-4.
56. Chen AF, Cao D-S, Xiao N, Xu Q-S. Rcpi: R/Bioconductor package to generate various descriptors of proteins, compounds and their interactions. Bioinformatics. 2014; 31:279-81.
57. Waterhouse A, Bertoni M, Bienert S, Studer G, Tauriello G, Gumienny R, et al. SWISS-MODEL: homology modelling of protein structures and complexes. Nucleic Acids Res. 2018; 46:W296-W303.
58. Trott O, Olson AJ. AutoDock Vina: improving the speed and accuracy of docking with a new scoring function, efficient optimization, and multithreading. Journal of Computational Chemistry. 2010; 31:455-61.
59. Morris GM, Huey R, Lindstrom W, Sanner MF, Belew RK, Goodsell DS, et al. AutoDock4 and AutoDockTools4: Automated docking with selective receptor flexibility. Journal of Computational Chemistry. 2009; 30:2785-91.
60. Pettersen EF, Goddard TD, Huang CC, Couch GS, Greenblatt DM, Meng EC, et al. UCSF Chimera—A visualization system for exploratory research and analysis. Journal of Computational Chemistry. 2004; 25:1605-12.
61. Forli S, Huey R, Pique ME, Sanner MF, Goodsell DS, Olson AJ. Computational protein-ligand docking and virtual drug screening with the AutoDock suite. Nature protocols. 2016; 11:905-19.
62. Huey R, Morris GM, Forli S. Using AutoDock with AutoDockTools: a tutorial. 2005. https://autodock.scripps.edu/faqs-help/tutorial/using-autodock-with-autodocktools/UsingAutoDockWithADT_v2e.pdf
63. Huey R, Morris GM, Forli S. Using AutoDock 4 and AutoDock Vina with AutoDockTools: A Tutorial. 2012. https://autodock.scripps.edu/faqs-help/tutorial/using-autodock-4-with-autodocktools/2012ADTtut.pdf.
64. Yang H, Tan L, Sala R, Burley SK, Hudson BP, Bhikadiya C, et al. RCSB Protein Data Bank: biological macromolecular structures enabling research and education in fundamental biology, biomedicine, biotechnology and energy. Nucleic Acids Res. 2018; 47:D464-D74.
65. Gindulyte A, Shoemaker BA, Yu B, He J, Zhang J, Chen J, et al. PubChem 2019 update: improved access to chemical data. Nucleic Acids Res. 2018; 47:D1102-D9.
66. Okonechnikov K, Golosova O, Fursov M, the Ut. Unipro UGENE: a unified bioinformatics toolkit. Bioinformatics. 2012; 28:1166-7.
67. Altschul SF, Madden TL, Schaffer AA, Zhang J, Zhang Z, Miller W, et al. Gapped BLAST and PSI-BLAST: a new generation of protein database search programs. Nucleic Acids Res. 1997; 25.

68. Umeda T, Katsuki J, Ashikawa Y, Usami Y, Inoue K, Noguchi H, et al. Crystallization and preliminary X-ray diffraction studies of a terminal oxygenase of carbazole 1,9a-dioxygenase from Novosphingobium sp. KA1. Acta Crystallographica Section F. 2010; 66:1480-3.
69. Parada AE, Needham DM, Fuhrman JA. Every base matters: assessing small subunit rRNA primers for marine microbiomes with mock communities, time series and global field samples. Environ Microbiol. 2016; 18:1403-14.
70. Apprill A, McNally S, Parsons R, Weber L. Minor revision to V4 region SSU rRNA 806R gene primer greatly increases detection of SAR11 bacterioplankton. Aquat Microb Ecol. 2015; 75:129-37.
